# Arrestin-3 scaffolds multiple MAP3Ks driving stress-induced JNK3 activation and cell death

**DOI:** 10.64898/2026.03.09.710604

**Authors:** Chen Zheng, Mohamed R. Ahmed, Richard Sando, Vsevolod V. Gurevich, Eugenia V. Gurevich

**Author notes:** **Correspondence:** Eugenia V. Gurevich, and Vsevolod V. Gurevich,; Department of Pharmacology, Vanderbilt University, Nashville, TN 37232.

## Abstract

Non-visual arrestin-3 (a.k.a. β-arrestin-2) functions as a scaffold facilitating the activation of c-Jun N-terminal kinases (JNKs), an important pathway regulating cell fate. Here, we demonstrate that arrestin-3 scaffolds not only previously identified ASK1, but facilitates signaling by several MAP3Ks, including ZAKα, ZAKβ, MEKK1, and TAK1. We identified ZAK (sterile alpha motif and leucine zipper-containing kinase) as the predominant MAP3K mediating arrestin-3-dependent JNK3 signaling and chemotherapy drug-induced cell death in HEK293 cells. We also showed that a 16-residue-long arrestin-3-derived peptide binds ZAK and fulfills the scaffolding function of full-length arrestin-3, sensitizing cells to death induced by chemotherapy drugs. These findings demonstrate that arrestin-3 is a versatile facilitator of stress signaling and suggest that functional peptide mimics can be used therapeutically to facilitate drug-induced death of cancer cells.

## Introduction

Arrestins were originally discovered as negative regulators of G protein-mediated signaling of G protein-coupled receptors (GPCRs) (reviewed in (Carman and Benovic, 1998)). Both non-visual arrestins (arrestin-2 and arrestin-3)^1^ were also shown to act as signal transducers organizing multiprotein complexes (reviewed in (Peterson and Luttrell, 2017; Wess et al., 2023; Gurevich and Gurevich, 2024)). The role of arrestins in the signaling via the mitogen-activated kinase (MAPK) pathways was extensively studied (McDonald et al., 2000; Luttrell et al., 2001; Bruchas et al., 2006; Wess et al., 2023; Ahmed et al., 2024). MAPKs regulate vital cellular functions: survival, proliferation, differentiation, migration, and death (Pearson et al., 2001; Johnson and Lapadat, 2002; Johnson, 2011; Peterson et al., 2022). MAPK activation cascades are composed of kinases of three levels; the upstream kinases sequentially activate the downstream kinases by phosphorylation. The downstream effector MAPKs (fourteen in humans, including ERK1/2, JNKs, p38) are doubly phosphorylated by intermediate MAPK kinases (MAP2Ks; seven in humans), whereas these are phosphorylated by the upstream MAPK kinase kinases (MAP3Ks; twenty-four different ones in humans) (Manning et al., 2002; Peterson et al., 2022).

The mode of activation of MAP3Ks is complex including phosphorylation or dephosphorylation of select residues by other protein kinases and phosphatases, autophosphorylation, binding to various regulatory proteins and membrane, as well as dimerization in response to diverse extracellular stimuli and intracellular conditions (Xia et al., 2000; Sakurai, 2012; Xu and Lei, 2020; Mordente et al., 2024) thus conferring specificity to the cell response (Craig et al., 2008; Peterson et al., 2022). Different MAP3Ks respond to distinct signals, some of which are pro-survival, others indicate various types of stress (Xia et al., 2000; Craig et al., 2008; Xu and Lei, 2020; Snieckute et al., 2022; Mordente et al., 2024). MAP3Ks ultimately control the activity of the MAPK signaling pathways and the cellular response is determined by the activation of specific MAP3Ks via the coordinated activity of the MAPKs (Johnson et al., 2005; Craig et al., 2008; Johnson, 2011; Peterson et al., 2022; Mordente et al., 2024).

The MAPK pathways often rely on scaffold proteins that facilitate signaling in the three-kinase cascades as well as confer spatial specificity and reduce crosstalk (Brown and Sacks, 2009; Musi et al., 2020). Arrestins have been shown to act as scaffolds to the MAPK pathways (McDonald et al., 2000; Luttrell et al., 2001; Bruchas et al., 2006). Arrestin-3 (a.k.a. β-arrestin2) is unique in the family in its ability to facilitate the activation of the c-Jun N-terminal kinases (JNKs), with particular preference for the neuronal isoform JNK3 (McDonald et al., 2000; Song et al., 2009; Zhan et al., 2011b; Zhan et al., 2013; Kook et al., 2014; Zhan et al., 2023), by serving as a scaffold of the JNK activation module. This function is important biologically because JNK signaling is critical for the stress responses and cell death or survival (Yan et al., 2024; Vind et al., 2025). Specifically, the JNK3 signaling governs the balance between neuronal survival and death in response to environmental stress and neurotoxic insults (Bonny et al., 2005; Castro-Torres et al., 2024; Mordente et al., 2024). Dysregulation of the JNK3 signaling was implicated in neurodegeneration (Antoniou et al., 2011; Musi et al., 2020; de Los Reyes Corrales et al., 2021; Zhao et al., 2022), making the mechanism of the assembly of JNK3-activating signaling cascades a subject of intense investigation.

Arrestin-3 was previously reported to facilitate the signaling of only one MAP3K upstream of JNK3, apoptosis signal-regulated kinase 1 (ASK1) (McDonald et al., 2000), which responds to oxidative stress (Guo et al., 2017), via scaffolding of the ASK1-MKK4/MKK7-JNK3 module. However, this model does not explain observed facilitation of JNK3 activation by arrestin-3 in the absence of ASK1 (Perry-Hauser et al., 2022). Therefore, we tested whether arrestin-3 facilitates signal transduction from other MAP3Ks to JNK3. We found that arrestin-3 facilitates JNK3 activation initiated by multiple MAP3Ks, including ZAKα, ZAKβ, MEKK1/2/4, and TAK1. These MAP3Ks are activated by different stressors: TAK1 (transforming growth factor-β-activated kinase 1) responds to cytokines, ZAKα (sterile alpha motif and leucine zipper-containing kinase) to ribotoxic stress, ZAKβ, a shorter splice variant of ZAKα, to mechanical stress, MEKK1 to osmotic stress, etc. (reviewed in (Mordente et al., 2024)). We identified the two isoforms of endogenous ZAK as strong drivers of arrestin-3-facilitated JNK3 activation in HEK293 cells, where ASK1 plays a negligible role. We showed that a short 16-residue peptide derived from the N-terminus of arrestin-3 (T16) also facilitates JNK3 activation initiated by several MAP3Ks, including ZAK, and mimics the pro-death function of the full-length arrestin-3 in HEK293 cells. These findings define arrestin-3 as a versatile scaffold that facilitates signal transduction from many MAP3Ks to enhance stress responses and demonstrate that arrestin-3-derived mini-scaffold peptides are usable tools for the regulation of cell fate.

## Results

### Arrestin-3 scaffolds diverse MAP3Ks

First, we tested arrestin-3 interactions with a panel of seven MAP3Ks representing structurally diverse families (Peterson et al., 2022; Mordente et al., 2024). Co-immunoprecipitation was performed in HEK293 arrestin-2/3 knockout (AKO) cells to preclude the interference of endogenous arrestins. We found that arrestin-3 interacts with every MAP3K tested, including ASK1, ZAK (both α and β isoforms), MEKK1, MEKK2, MEKK4, and TAK1 (**Fig. 1A, B**). While the affinities varied, with ASK1 and ZAKβ showing the strongest binding, these data showed that arrestin-3 interacts with a variety of MAP3Ks.

**Figure 1.**
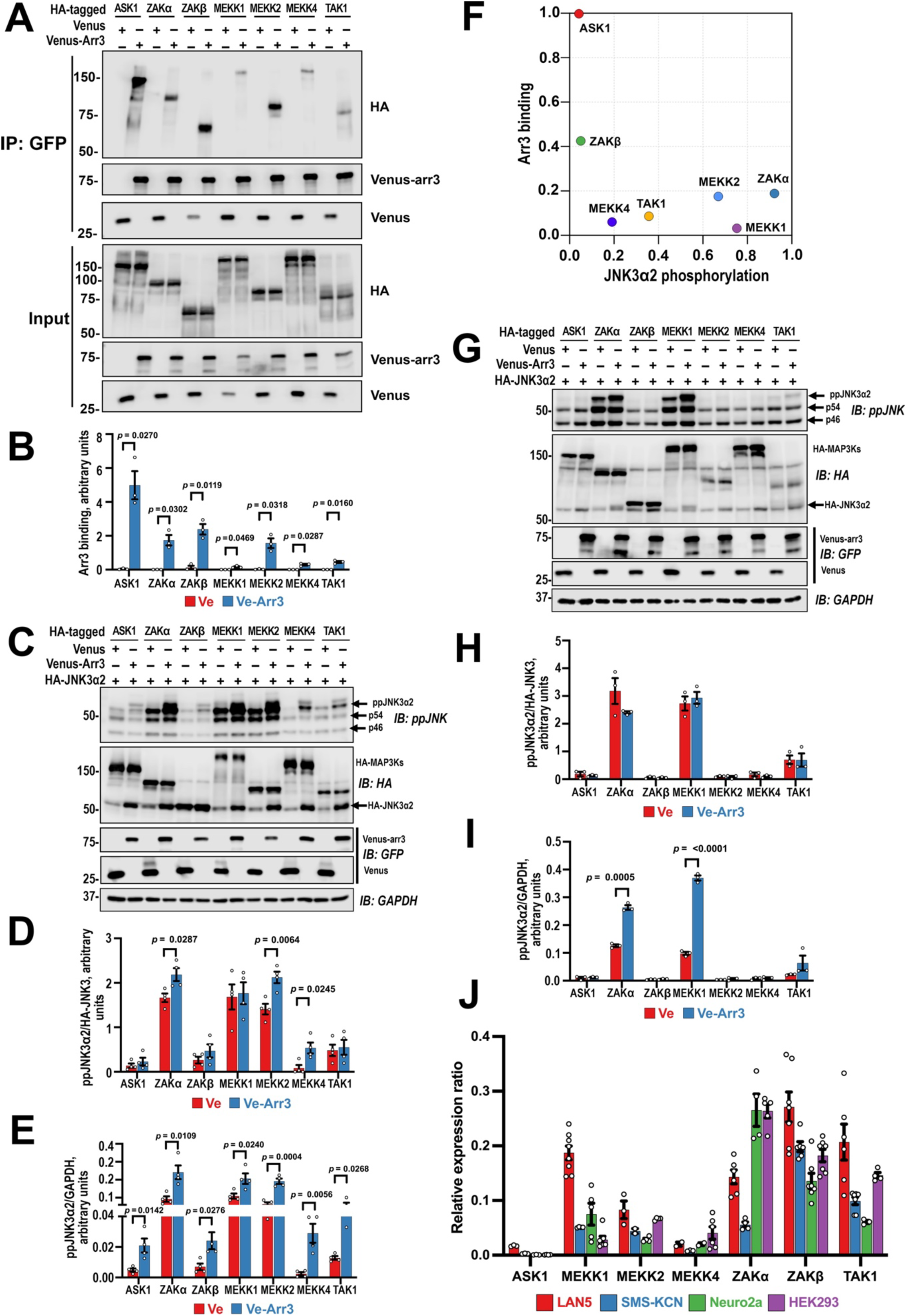
Arrestin-3 binds several MAP3Ks and facilitates JNK3 activation triggered by these kinases. **A**. AKO cells were co-transfected with Venus-arrestin-3 and indicated HA-MAP3Ks. Venus-arrestin-3 was precipitated using GFP-trap beads and bound HA-MAP3Ks were detected by western blotting. **B.** Quantification of **A** (N=3). **C.** Arrestin-3 effect on JNK3 activation driven by different MAP3Ks in AKO cells co-transfected with HA-JNK3α2, indicated HA-MAP3Ks, and Venus (Ve) or Venus-arrestin-3 (Ve-arr3). JNK3 activation was determined by immunoblotting for doubly phosphorylated JNK (ppJNK). **D, E.** Quantification of **C.** Bar graphs show JNK3 activation normalized to total HA-JNK3α2 (**D**) or the loading control GAPDH (**E**) (means +/− SEM; N=3). **F.** MAP3K binding vs arrestin-3-dependent increase in JNK3 activation in AKO cells. **G.** Arrestin-3 effect on JNK3 activation driven by different MAP3Ks in Neuro2a cells. Experiments were performed as in **C. H, I.** Quantification of **G** normalized to total HA-JNK3α2 (**H**) or GAPDH (**I**). **J.** Relative expression of endogenous MAP3Ks in different cell lines (HEK293, Neuro2a, LAN5, SMS-KCN) (quantification strategy is shown in Figs. S3, S4). For panels **B, D, E, H**, and **I**, statistical significance of the differences between Ve and Ve-Arr3 within each specific MAP3K transfection group was determined using multiple unpaired t-tests with Welch’s correction, and the False Discovery Rate (FDR) was set at 5%; p values are indicated.

We next determined whether these interactions translate into facilitation of JNK3 activation in two distinct cellular contexts: mouse neuroblastoma Neuro2a and HEK293-derived AKO cells. As many MAP3Ks were shown to dimerize and self-activate at high expression levels (Blank et al., 1996; Sakurai et al., 2000; Cheng et al., 2005b; Cheng et al., 2005a; Nishida et al., 2017; Pleinis et al., 2017; Köster et al., 2024; Vish et al., 2025; Huso et al., 2026), we used overexpression as a means of activating MAP3Ks. MAP3Ks differentially affected phosphorylation levels of expressed JNK3α2 and endogenous JNK isoforms p54 and p46, with MEKK1, ZAKα, MEKK2, and ASK1 being the most, while ZAKβ and MEKK4 the least effective in all cases (**Fig. S1**). TAK1 showed higher relative effectiveness towards p46 than towards p54 and JNK3α2 (Fig. S1). It is likely that varying levels of the activation of different isoforms of JNK observed, in addition to possible MAP3K specificity, are determined by variable ability of different MAP3Ks to self-activate upon overexpression and/or use other scaffolds endogenously expressed by AKO cells. Because we were exclusively interested in the ability of arrestin-3 to facilitate JNK3α2 activation initiated by each MAP3K, this does not affect the conclusions based on the data in Fig. 1. On the AKO background, arrestin-3 potentiated JNK3 activation driven by all tested MAP3Ks (**Fig. 1C, E**). In contrast, in Neuro2a cells, the arrestin-3 effect was more selective: it enhanced signaling driven by ZAKα and MEKK1 with minimal effect on MEKK2, MEKK4, TAK1, and ASK1 (**Fig. 1G, I**).

We consistently found that arrestin-3 co-expression increased the level of expressed HA-JNK3α2 (**Fig. 1C, G, middle panels**). We have examined this effect directly by expressing increasing levels of Venus-arrestin-3 in AKO cells and measuring the corresponding co-expression of HA-JNK3α2 driven by constant amount of HA-JNK3α2 encoding plasmid (**Fig. S2A, B**). We found that the expression of HA-JNK3α2 rose with the co-expression of Venus-arrestin-3, with covariation coefficient ∼0.98 (Fig. S2C). In contrast, Venus-arrestin-3 had no effect on the expression levels of endogenous MAP2Ks known to phosphorylate JNKs, MKK4 and MKK7 (**Fig. S2A, D, E**). These data suggest that a fraction of free arrestin-3 in the cytoplasm exists in pre-assembled complexes with JNK3α2, but likely not with MKK4/7.

**Fig. 1D, H** shows the level of ppJNK3 normalized to the total amount of HA-JNK3α2, which reflects the contribution of arrestin-3 specifically to the activation of expressed JNK3α2 via increased phosphorylation. As the biologically relevant signal the cell receives is proportional to the absolute number of active JNK3α2 molecules, to rigorously quantify the strength of the signal in the presence of arrestin-3, we normalized the amount of doubly phosphorylated (fully active) JNK3α2 to the loading control (GAPDH) (**Fig. 1E, I**). These panels show total increase in the strength of the signal (the number of active JNK3α2 molecules) caused by arrestin-3 in the presence of different MAP3Ks, as compared to the level of ppJNK3α2 activated by the same MAP3Ks in the absence of arrestin-3.

In AKO cells, the strength of the binding of individual MAP3Ks to arrestin-3, as measured by co-immunoprecipitation, did not correlate with its ability to enhance JNK3 activation driven by these MAP3Ks (**Fig. 1F**). For example, MEKK1 bound arrestin-3 weakly (**Fig. 1A, B**) but arrestin-3 robustly increased JNK3 activation induced by MEKK1 (**Fig. 1C, D, E**), suggesting that transient interactions are sufficient for efficient scaffolding. Collectively, these data indicate that arrestin-3 is not ASK1-specific, but functions as a versatile scaffold of JNK3-activating cascades initiated by several stress-responsive MAP3Ks.

### Endogenous MAP3Ks in different cell types

As different cells express distinct complements of MAP3Ks, the versatility of arrestin-3 as the scaffold of JNK3-activating cascades suggests that its biological function is likely cell subtype-specific. MAP3Ks have been reported to act in concert in response to various stimuli thereby determining cell fate via unique combination of the activity of individual MAP3K (Chen et al., 2002; Peterson et al., 2022). Cells usually express several MAP3K subtypes, and it stands to reason that the cellular complement of MAP3Ks, particularly the relative abundance of individual MAP3Ks in each cell type, would impact signaling responses. However, the information regarding the expression of endogenous MAP3Ks in different cells remains scarce.

The quantification of MAP3K expression in absolute values is impossible, since purified individual MAP3K proteins necessary to construct calibration curves are not available. Therefore, we used a two-step quantitative immunoblotting to determine the relative abundance of select MAP3Ks in wild type HEK293 (human embryonic kidney) cells, Neuro2a (mouse neuroblastoma), and two human neuroblastoma lines, LAN5 and SMS-KCN (**Fig. S3A**). First, calibration curves were established using increasing amounts of transfected HA-tagged MAP3Ks detected by anti-HA antibody, allowing us to equalize the relative amounts of distinct MAP3K standards (**Fig. S3B**). Next, MAP3K-specific antibodies were used to detect both the endogenous MAP3Ks and the pre-quantified HA-tagged MAP3Ks serving as reference standards on the same blots. The standard curves generated from the HA-references were then utilized to determine the relative abundance of endogenous kinases in whole-cell lysates (this procedure for ZAKα is shown in Fig. S3C, for other MAP3Ks in Fig. S4). This strategy allowed an accurate quantification of the relative abundance of endogenous MAP3Ks.

We found that the complement of endogenously expressed MAP3Ks in these cell types is different (**Fig. 1J**). In AKO cells expression of ZAK (both α and β isoforms) and TAK1 was significantly greater than that of ASK1 and MEKK1/2/4; in Neuro2a cells ZAKα was the most abundant, followed by ZAKβ and MEKK1; in SMS-KCN cells ZAKβ and MEKK1 were the most abundant MAP3Ks; in LAN5 cells MEKK1 was the highest expressed MAP3K, followed by ZAKβ (**Figs. 1J, S4**). Endogenous expression of ASK1, previously shown to interact with arrestin-3 (McDonald et al., 2000), as well as of MEKK2 and MEKK4, was relatively low in all cell types tested (**Figs. 1J, S4**).

### The role of ZAK in arrestin-3-dependent JNK3 activation in AKO cells

Because of high abundance of ZAK (**Figs. 1J, S4**) and its robust responsiveness to arrestin-3 scaffolding in AKO cells (**Fig. 1C, E**), we hypothesized that ZAK might be the primary player in the arrestin-3-dependent facilitation of JNK3 activation in this cell type. To test this hypothesis, we generated ZAK knockout (KO) lines on the background of AKO cells (thereby creating triple-null cell lines) using CRISPR/Cas9 (**Fig. 2A**). We validated the knockout of ZAK gene and complete loss of ZAK proteins in two independent clones (**Fig. S5**). The loss of ZAK significantly reduced the basal JNK3 activity and abolished most of arrestin-3-mediated JNK3 activation in AKO cells (**Fig. 2B, C**). This deficiency was observed with full-length arrestin-3 and two arrestin-3-derived peptides (T16 and T14) previously shown to facilitate JNK3 activation (Perry-Hauser et al., 2022). Thus, in this cellular context all three scaffolds operate primarily by assisting endogenous ZAK.

**Figure 2.**
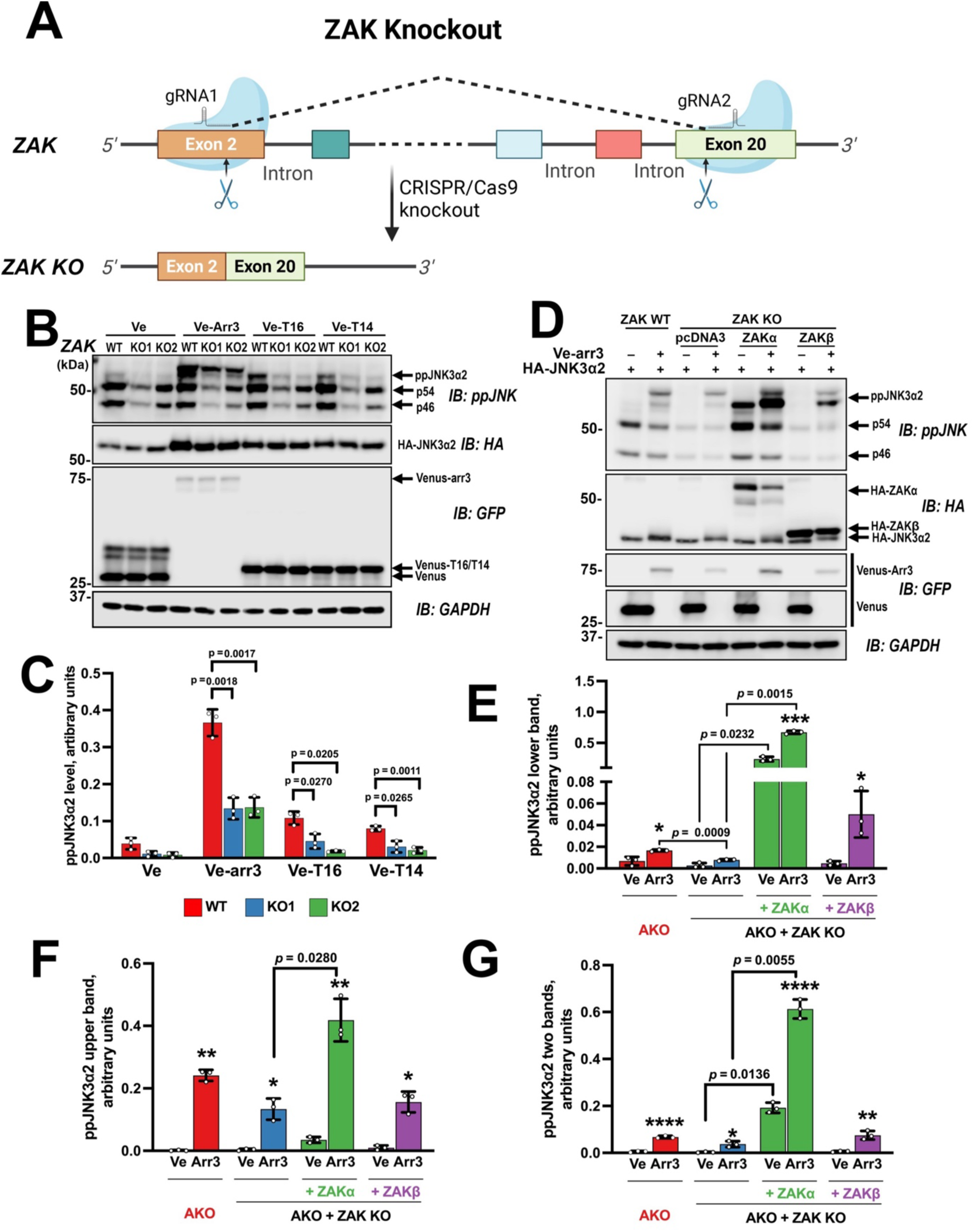
ZAK mediates most arrestin-3 effects on JNK3 activation in HEK293 cells. **A**. Schematic of the CRISPR/Cas9-mediated ZAK knockout strategy targeting exons 2 and 20. **B.** The effect of ZAK knockout on arrestin-3– and peptide-facilitated JNK3 activation. Parental (AKO) and two independent ZAK knockout clones (KO1, KO2) were transfected with HA-JNK3α2 and Venus (Ve), Venus-arrestin-3 (Ve-Arr3), Venus-T16 (Ve-T16), or Venus-T14 (Ve-T14). Peptide sequences are shown in Fig. 3B. Western blots and PCR analysis of AKO cells and knockout clones are shown in Fig. S5. **C.** Quantification of JNK3α2 phosphorylation in **B**. JNK3 activation (both bands together) was assessed by immunoblotting with anti-phospho-JNK antibody. **D.** AKO or AKO + ZAK KO cells (clone KO1) transfected pcDNA3 (control), HA-ZAKα, or HA-ZAKβ and co-transfected with HA-JNK3α2 and Venus (control; Ve) or Venus-arrestin-3 (Ve-Arr3). Immunoblotting revealed two distinct ppJNK3α2 bands: a lower band (ZAK-dependent) and an upper band (ZAK-independent). Quantification of lower band (**E**), upper band (**F**), and both ppJNK3α2 bands together (**G**) (means ± SEM; N=3) in **(D)**, statistical significance was determined by Welch’s one-way ANOVA followed by Dunnett’s correction for multiple comparisons. For panels **(E-G)**, comparisons between Venus (Ve) and Venus-arrestin-3 (Ve-Arr3) within each specific genotype were analyzed using multiple unpaired t-tests with FDR set at 5%. To evaluate the effect of a given treatment (Ve or Ve-Arr3) on different genetic backgrounds in **(E-G)**, Welch’s one-way ANOVA was performed, followed by Dunnett’s multiple comparisons test. P-values are indicated.

We confirmed the specificity of this effect by rescue experiments. The expression of ZAKα and to a lesser extent ZAKβ in ZAK KO cells restored arrestin-3 facilitation of JNK3α2 activation (**Fig. 2D**). Immunoblots revealed that activated doubly phosphorylated JNK3α2 (ppJNK) runs as two distinct molecular species: a “lower” band and an “upper” band (**Fig. 2D**). The lower band is strictly ZAK-dependent, disappearing in ZAK KO cells and reappearing upon the expression of either isoform of ZAK (Fig. 2D, E). The upper band, which is also enhanced by the expression of arrestin-3, exhibits partial resistance to ZAK knockout (Fig. 2D, F), suggesting that it likely represents the population of JNK3α2 activated by other endogenous MAP3Ks (e.g., TAK1, which is also abundant in these cells) with the help of the arrestin-3 scaffold (**Fig. 1**). The sum of both bands (Fig. 2G) shows the integral action of arrestin-3 on JNK3α2 activation. The profound reduction of arrestin-3-facilitated JNK3α2 activation in ZAK KO cells demonstrated that ZAK is the most important partner of arrestin-3 in AKO and likely parental HEK293 cells.

### Peptide mimics retain the scaffolding function of arrestin-3

Next, we tested whether short JNK3-activating peptides derived from the arrestin-3 N-terminus (Perry-Hauser et al., 2022) facilitate signaling of different MAP3Ks to JNK3 like the full-length protein. The advantage of these peptides as experimental tools is their limited functionality as compared to the remarkably multi-functional full-length arrestin-3 (Xiao et al., 2007; Gurevich and Gurevich, 2019). We tested whether arrestin-3-derived peptides T1A (residues 1-25), T16 (residues 9-24), T15 (residues 7-21), and T14 (residues 11-24) (**Fig. 3A, B**) facilitate MAP3K-driven JNK3 activation in Neuro2a cells. We found that the ability of the peptides to facilitate signaling of different MAP3Ks varied: T1A and T14 were more potent than full-length arrestin-3 with MEKK2, T16 with ASK1, MEKK4, and ZAKβ, whereas T15 acted like arrestin-3 with TAK1, but was less effective with other MAP3Ks (**Figs. 3C, S7**). Thus, the peptides representing different parts of the arrestin-3 N-terminus are more selective towards individual MAP3Ks than full-length protein. Observed difference in peptide activity is determined by their ability to engage MAP3Ks, as all downstream cascade components are the same (MKK4/7 and JNK3α2). All tested peptides share eleven residues but differ in N– and C-termini (Fig. 3B). Based on their ability to assist tested MAP3Ks, the N-terminal six residues and/or the C-terminal Arg of T1A are inhibitory, whereas the C-terminal Tyr-Leu-Gly (present in T16 and T14, absent in T15) preferentially improve their activity with ASK1 and MEKK4. Overall, T16 was the most consistent functional mimic of full-length arrestin-3, robustly enhancing JNK3α2 activation driven by multiple MAP3Ks (**Fig. 3C**; western blots and quantification shown in **Figs. S6, S7**). Thus, T16 is effective in cells of neuronal origin that endogenously express JNK3 isoform.

**Figure 3.**
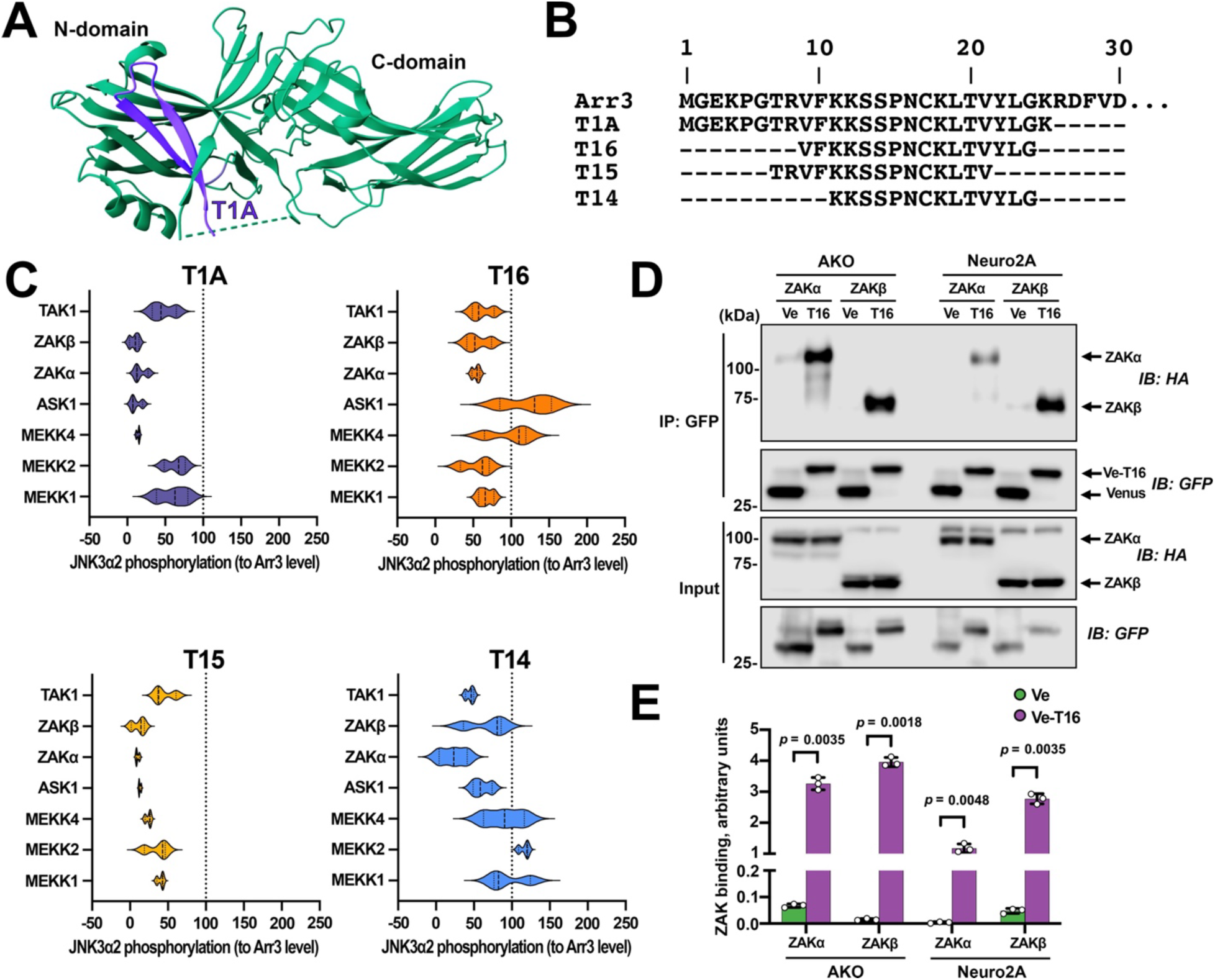
Arrestin-3-derived peptides mimic its scaffolding function in Neuro2a cells. **A**. Structure of arrestin-3 (PDB ID 3P2D) with the sequence corresponding to the N-terminal T1A peptide shown in blue. **B.** Amino acid sequences of arrestin-3 (Arr3) and derived peptides. **C.** Neuro2a cells were transfected with HA-JNK3α2, indicated MAP3Ks, and Venus-tagged peptides or Venus (control). Violin plots show the distribution of JNK3α2 activation effects (shown as % of full-length arrestin-3 efficacy indicated as vertical dotted lines). Blots and quantification are in **Figs. S6, S7**. **D.** Co-immunoprecipitation was performed from AKO cells transfected with Venus (Ve) or Venus-T16 (Ve-T16) and HA-ZAKα or HA-ZAKβ. **E.** Quantification of the binding of Venus-T16 to ZAKα and ZAKβ. Data represent means ± SEM (N= 3). Statistical significance was determined by multiple unpaired t-tests with FDR set at 5%; p-values are indicated above bars.

To test the interaction of T16 with ZAK, we co-expressed Venus-T16 with HA-ZAKα and HA-ZAKβ in AKO and Neuro2a cells, immunoprecipitated Venus-T16 with GFP trap beads, and tested for the presence of HA-ZAK isoforms in the precipitate. We found that T16 interacts with both ZAKα and ZAKβ (**Fig. 3D, E**). To further validate the utility of peptide tools, we similarly tested the shortest peptide T14 (residues 11-24) and found that T14 also binds MAP3Ks tested, TAK1 and ZAKβ (**Fig. S8**). These data are consistent with the idea that arrestin-3-derived peptides facilitate JNK3 activation by functioning as mini-scaffolds that assemble productive three-tier MAP3K-MAP2K-MAPK modules.

### Arrestin-3 potentiates JNK3 activation induced by stressors

To evaluate the biological role of arrestin-3 scaffolding we tested JNK3α2 activation under stress conditions using compounds known to activate MAPK pathways: anisomycin, a protein synthesis inhibitor and potent activator of the JNK pathway (Mahadevan and Edwards, 1991; Curtin and Cotter, 2002; Rosser et al., 2004), and two chemotherapy drugs vincristine and doxorubicin (Fig. 4). Anisomycin triggers ribotoxic stress response (RSR), which is mediated by ZAKα (Vind et al., 2020; Vind et al., 2025). Doxorubicin, a DNA damage-inducing chemotherapy agent, has been also shown to activate RSR via ZAK (Sauter et al., 2010). Vincristine, a microtubule destabilizer, has been shown to activate the JNK in sensory neurons and carcinoma cells, which resulted in cell death and neurodegeneration (Stone and Chambers, 2000; Brantley-Finley et al., 2003; Tsai et al., 2022).

**Figure 4.**
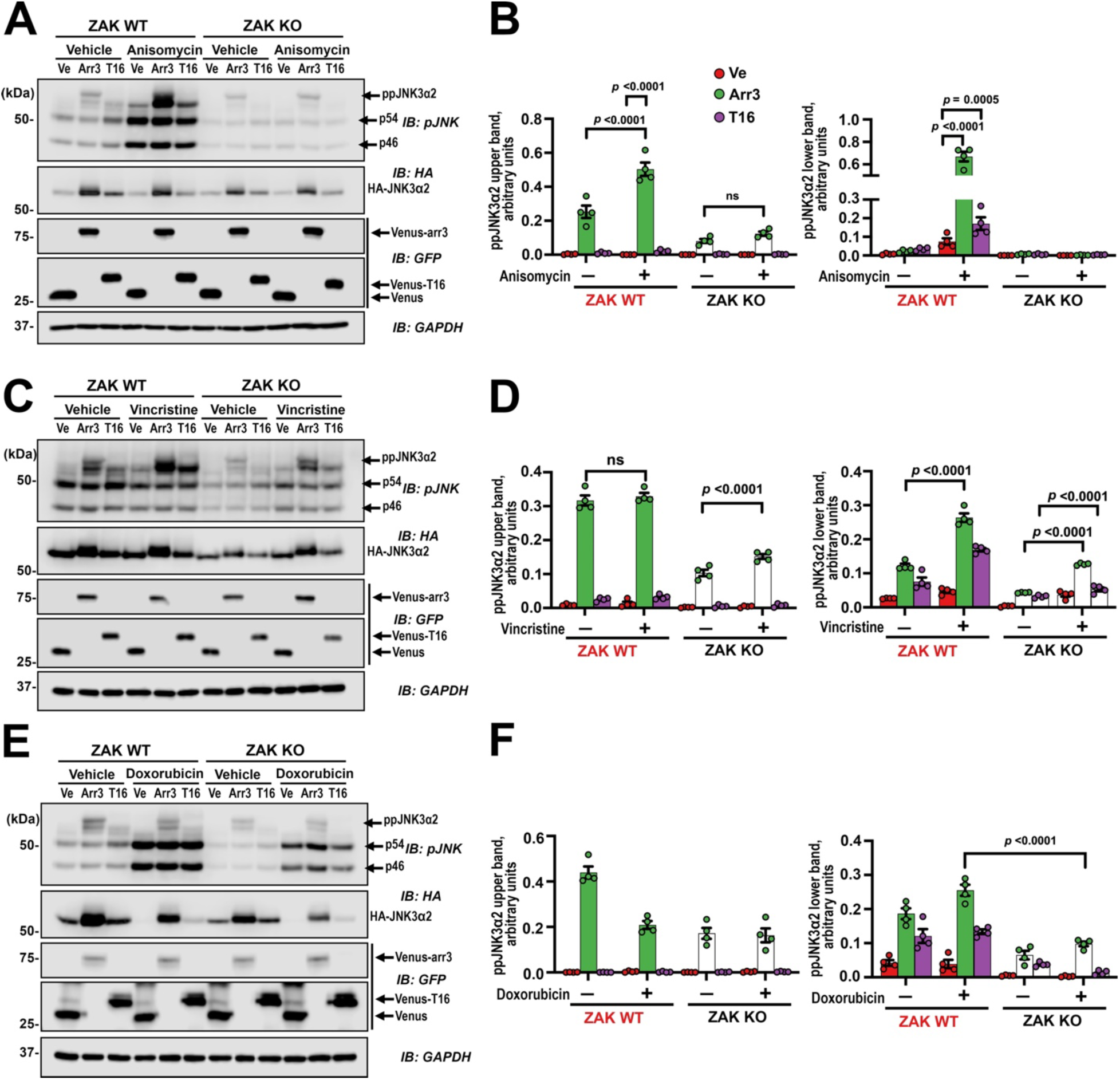
Dependence of arrestin-3 enhancement of stress-induced JNK3 activation on ZAK. **A, C, E.** Western blot for ppJNK3α2 in ZAK WT and ZAK KO AKO cells treated with anisomycin (300 nM, 1 h) (**A**), vincristine (100 nM, 24 h) (**C**), or doxorubicin (20 nM, 24 h)(**E**). Cells were transfected with HA-JNK3α2 and Venus (control; red), Venus-arrestin-3 (green), or Venus-T16 (magenta). Note the separation of ppJNK3α2 into ZAK-dependent (lower) and ZAK-independent (upper) bands. **B, D, F.** Quantification of ppJNK3α2 in panels **A**, **C**, and **E**, respectively. The upper and lower ppJNK3α2 bands were quantified separately, the intensity of upper (ZAK-independent) and lower (ZAK-dependent) bands is shown. Bar graphs represent means ± SEM (N=3). Statistical significance was determined by two-way ANOVA with Sidak’s correction for multiple comparisons; p values are indicated. Quantification of phosphorylation of endogenous p46 and p54 is shown in Fig. S9.

We found that short-term treatment with anisomycin (300 nM, 1 h) resulted in significant activation of JNK3α2, which was strictly ZAK-dependent (**Fig. 4A, B**). In AKO cells expressing endogenous ZAK, arrestin-3 and T16 peptide robustly enhanced the intensity of the lower ppJNK3α2 band, but this effect was completely abolished in ZAK KO cells (**Fig. 4A, B**). Thus, ZAK is the main transducer for anisomycin-induced JNK3 activation in AKO cells.

Prolonged microtubule stress (vincristine, 100 nM, 24 h) revealed a more nuanced regulation. While ZAK deletion significantly reduced arrestin-3– and T16-dependent enhancement of JNK3α2 activation (the lower ppJNK3α2 band in **Fig. 4C, D**), both bands remained responsive to arrestin-3 and T16 even in the absence of ZAK (**Fig. 4C, D**). Thus, while ZAK contributes significantly to vincristine-induced stress signaling, arrestin-3 and T16 also scaffold other MAP3Ks partially bypassing ZAK deficiency.

The response to genotoxic stress (doxorubicin, 20 nM, 24 h) was the most complex (**Fig. 4E, F**). Doxorubicin treatment led to a marked reduction in total HA-JNK3α2 expression in control and T16-expressing cells, whereas the expression of full-length arrestin-3 protected HA-JNK3α2. In doxorubicin-treated cells, arrestin-3 increased ppJNK3α2 levels in a largely ZAK-dependent manner. Notably, we also observed increased intensity of lower molecular weight phosphorylated JNK species (eight endogenous JNK1/2 isoforms that run as two bands, p46 and p54), which were similarly dependent on the expression of ZAK, suggesting that under genotoxic stress arrestin-3 facilitates ZAK-dependent activation of several isoforms of JNK (Fig. S9).

### Stress signals trigger translocation of ZAKα and arrestin-3 to the same cytoplasmic regions

To visualize possible stress-induced movements of signaling proteins involved in the response, we analyzed intracellular distribution of arrestin-3 and ZAKα under basal and stress conditions. In control cells both Venus-arrestin-3 (green) and mCherry-ZAKα (red) exhibited diffuse distribution throughout the cytoplasm. Both proteins appeared to be excluded from the nucleus. Co-expressed Venus-arrestin-3 and mCherry-ZAKα showed the same diffuse localization (Fig. **5A**, third row). Line scans in co-expressing cells showed overlapping fluorescence profiles, but no subcellular locations where the proteins might be concentrated were detected (**Fig. 5B**).

**Figure 5.**
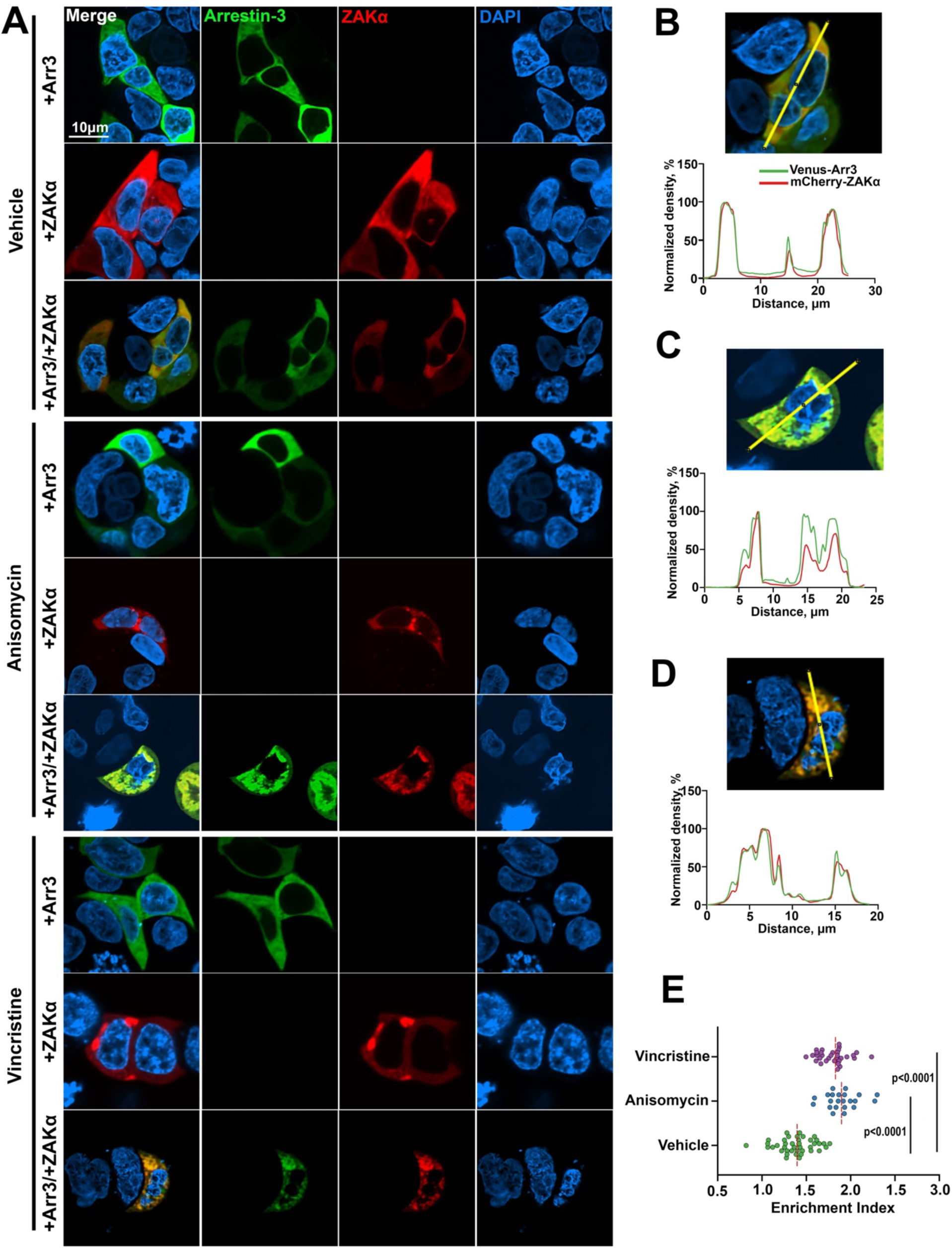
Arrestin-3 and ZAKα relocalize upon stress stimulation. **A**. Representative confocal images of AKO cells expressing Venus-arrestin-3 (green) or mCherry-ZAKα (red) alone or co-expressing both proteins. Cells were treated with vehicle (DMSO control), anisomycin (300 nM, 16h), or vincristine (500 nM, 16h). Scale bar: 10 μm. Nuclei are stained with DAPI (blue). Insets show magnified views of the individual co-expressing cells. **B, C, D.** Normalized fluorescence intensity line scans corresponding to the yellow lines in the images for control (**B**), vincristine (**C**), and anisomycin (**D**) treated co-expressing cells. **E.** Quantification of colocalization using the Enrichment Index (ratio of mean intensity in puncta to mean intensity in the cell). Each dot (control, green; anisomycin, blue; vincristine, magenta) represents a single cell; statistical significance of the differences was determined using one-way ANOVA with Dunnett’s correction for multiple comparisons; p values are shown.

Stress stimulation by both anisomycin and vincristine triggered a significant spatial reorganization: mCherry-ZAKα became concentrated in distinct cytosolic areas (**Fig. 5A,** fifth and eighth rows). mCherry-ZAKα underwent the same redistribution in the presence of arrestin-3 (**Fig. 5**, sixth and ninth rows). Interestingly, Venus-arrestin-3 redistribution was not observed in the absence of co-expressed mCherry-ZAKα (Fig. 5A, fourth and seventh rows). Thus, stress stimuli drive redistribution of ZAKα, whereas redistribution of Venus-arrestin-3 depends on its interactions with ZAKα. These data suggest that at least a fraction of arrestin-3 is pre-bound to ZAKα, which allows this MAP3K to “drag” arrestin-3 with it. Line scan analysis of co-expressing cells confirmed the tight spatial correlation of fluorescence peaks in cells treated with anisomycin or vincristine (**Fig. 5C,D**). To rigorously quantify this transition from the basal to stress-induced distribution we calculated an Enrichment Index (Varandas et al., 2016; Ripoll et al., 2024). This metric revealed a highly significant accumulation of ZAKα and arrestin-3 in distinct areas of the cytoplasm upon treatment with either stressor (**Fig. 5E**). Thus, stress induces the accumulation of ZAKα and its scaffold in select areas of the cytoplasm. Increased concentrations of both in these regions likely facilitate efficient signal transduction.

### Enhancement of JNK3 signaling by arrestin-3 increases ZAK-dependent cell death

Finally, we tested whether arrestin-3-dependent amplification of stress signals (**Fig. 4**) affects cell survival. To this end, we measured stress-induced cell death using flow cytometry with LIVE/DEAD staining in AKO and triple-null (AKO with ZAK KO) cells (gating strategy is shown in Fig. S10). First, we investigated spontaneous (not induced by stressors) cell death. We observed progressive accumulation of dead AKO cells from 24h to 48h post-transfection with Venus (control) or Venus-arrestin-3 (where t=0 was the time of adding the transfection mixture to the cells) and this trajectory was significantly flattened in ZAK KO cells (**Fig. 6A, B**).

**Figure 6.**
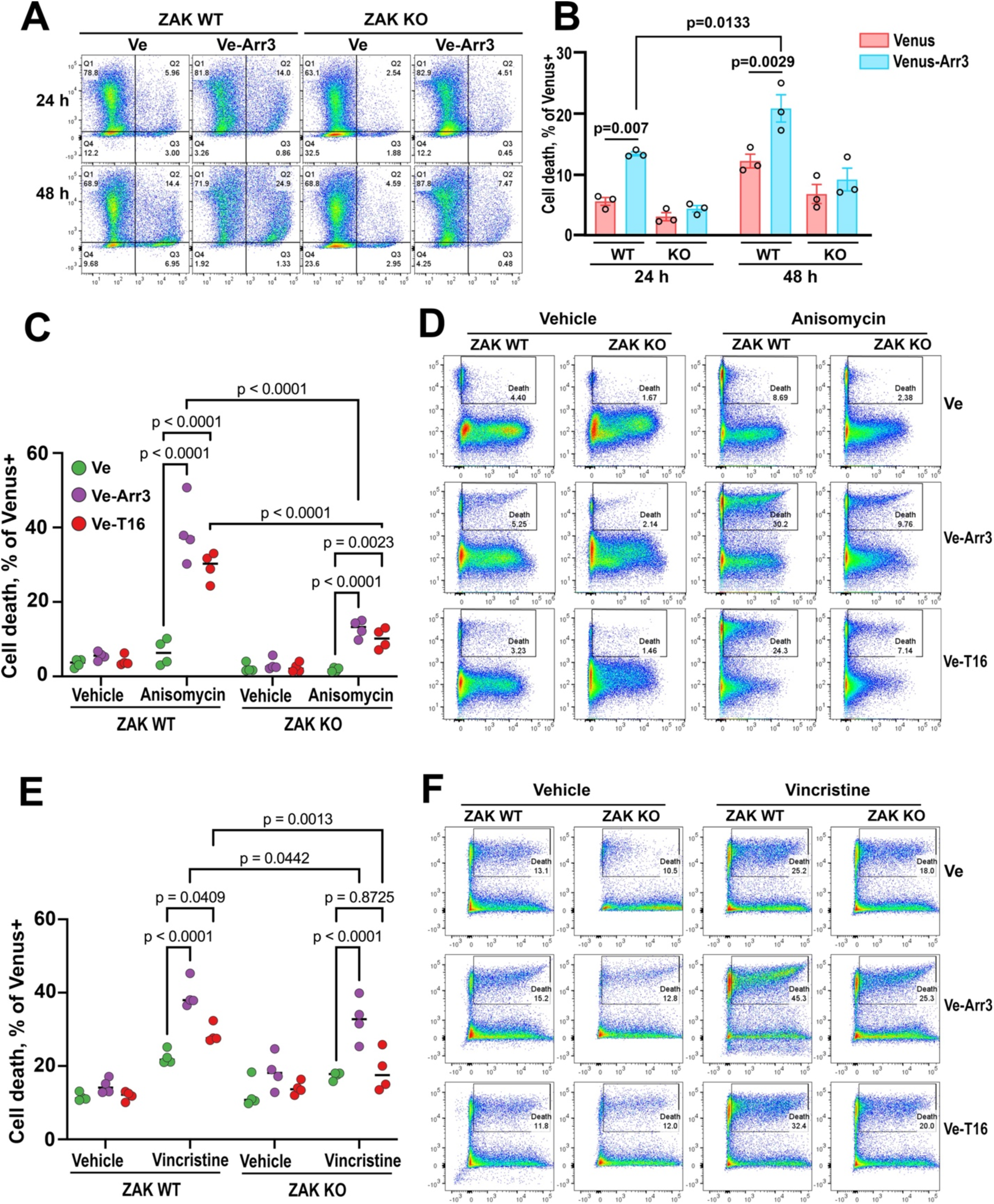
ZAK contribution to arrestin-3 and T16 effect on cell death. **A**. Time-dependent spontaneous cell death. Analysis of cell death in AKO (ZAK WT) and ZAK KO cells at 24h and 48h post-transfection with HA-JNK3α2 and Venus (control) or Venus-arrestin-3 in the absence of drug treatment. The time from transfection is indicated. Gating strategy is shown in Fig. S10. **B.** Quantification of panel **A.** Statistical significance was determined by three-way ANOVA with Sidak’s correction for multiple comparisons. p values are indicated. **C, D.** Anisomycin-induced cell death. **C**. Quantification of cell death of ZAK WT and ZAK KO AKO cells expressing Venus (control; green), Venus-arrestin-3 (magenta) or Venus-T16 (red) treated with DMSO (Vehicle) or anisomycin (1.5 μM, 48 h). **D.** Representative flow cytometry plots to panel **C**, gated for Venus+ populations. Numbers indicate the percentage of DEAD-stain positive cells. **E.** Vincristine-induced cell death. Quantification of cell death in cells treated with Vincristine (500 nM, 48 h). **F.** Representative flow cytometry plots corresponding to panel **E**. Bar graphs represent means ± SEM (N= 4). For panels **B**, **C**, and **E**, significance was determined by two-way ANOVA with Tukey’s correction for multiple comparisons; p values are indicated. The gating strategy and comprehensive flow cytometry analysis are shown in Figure S10.

Treatment with a lower dose of anisomycin (1.5 μM) induced a modest increase in death in control cells. The expression of arrestin-3 or the T16 in AKO cells sensitized them to anisomycin, as evidenced by a significant increase in the fraction of dead cells (**Fig. 6C, D;** detailed gating strategy and alternative visualization are provided in **Fig. S10**). This sensitization was strictly ZAK-dependent: in ZAK KO cells, neither arrestin-3 nor T16 increased cell death above control levels, effectively uncoupling arrestin-3 scaffold from the death machinery. Treatment with vincristine also increased cell death in AKO, and to a lesser extent in ZAK KO cells. This effect was enhanced by co-expression of arrestin-3 and T16 peptide (Fig. 6E, F). In this case the effect of arrestin-3 was much stronger than that of T16 (Fig. 6E).

Dose-response analysis revealed an intriguing functional difference between the full-length arrestin-3 and T16 peptide. In AKO cells, increasing anisomycin concentration to 3 μM further escalated cell death in the presence of arrestin-3. In contrast, T16 did not have this effect at the higher dose, suggesting that the full-length arrestin-3 possesses additional structural elements required for maximal facilitation of signaling under high-stress conditions (**Fig. S11**). The most likely reason for this is that the complexes organized by full-length arrestin-3 and T16 differ. It was shown that JNK3α2 binding is mediated by three distinct elements of arrestin-3 (Zhan et al., 2014), although only one of these, localized in the N-terminus, is capable of facilitating JNK3α2 activation in cells (Zhan et al., 2016; Perry-Hauser et al., 2022). Although the full set of the binding sites of MKK4, MKK7, and any MAP3K remains to be elucidated, it is likely that the other components of the activation cascade also engage elements of arrestin-3 absent in the N-terminal peptides. Importantly, in the absence of ZAK, this dose-dependency was completely lost for both arrestin-3 and T16 (Fig. S11).

Collectively, these data demonstrate that the arrestin-3/ZAK module acts as a critical rheostat for cellular susceptibility to stress, and that while T16 is a potent activator, the full scaffolding capacity of arrestin-3 provides the widest dynamic range of the pro-death signaling.

## Discussion

Three-tiered MAP3K-MAP2K-MAPK signaling modules are conserved in all eukaryotes, from yeast to mammals, as well as in plants (Widmann et al., 1999). Upstream MAP3Ks respond to a variety of stimuli and their activity determines which downstream effectors MAPKs are activated. Intricate interplay of the activity of different MAP3Ks regulates vital cellular functions and ultimately cell fate (Craig et al., 2008; Peterson et al., 2022). In particular, cellular responses to different types of stress are initiated by subsets of MAP3Ks (Craig et al., 2008; Peterson et al., 2022; Castro-Torres et al., 2024; Mordente et al., 2024). As MAPK activation cascades involve kinases of three levels, MAP3Ks, MAP2Ks, and MAPKs, signal transduction critically depends on scaffold proteins that bring sets of matching kinases together, thereby facilitating signaling (Dhanasekaran et al., 2007; Brown and Sacks, 2009).

While all arrestins share similar overall structure (Hirsch et al., 1999; Han et al., 2001; Sutton et al., 2005; Zhan et al., 2011a), the two non-visual subtypes demonstrate different functional capabilities and distinct tissue distribution (Wess et al., 2023; Gurevich and Gurevich, 2024). Non-visual arrestins were shown to function as scaffolds facilitating the activation of the three main families of MAPKs, ERK1/2 (Luttrell et al., 2001), JNKs (McDonald et al., 2000; Kook et al., 2014), and p38 (Bruchas et al., 2006). Both arrestin-2 and –3 scaffold ERK activation cascade only when they are bound to an activated and phosphorylated GPCR (Luttrell et al., 2001; Kolch, 2005; Breitman et al., 2012; Qu et al., 2021; Kim et al., 2022), whereas only arrestin-3 (but not arrestin-2) facilitates the activation of JNK family kinases (Miller et al., 2001; Song et al., 2009; Zhan et al., 2011b; Breitman et al., 2012; Zhan et al., 2013; Zhan et al., 2023). The JNK signaling was shown to be regulated by a family of scaffolding proteins, JIPs (JNK-interacting proteins) (Davis, 2000; Whitmarsh, 2006; Dhanasekaran et al., 2007). Arrestin-3 serves as an alternative scaffold for the JNK pathways with some preference for the JNK3 (Song et al., 2009; Zhan et al., 2011b; Zhan et al., 2013; Zhan et al., 2023).

Arrestin-3 was previously shown to scaffold JNK3-activating cascade initiated by only one stress-responsive MAP3K, ASK1 (McDonald et al., 2000; Song et al., 2009; Zhan et al., 2016). This suggested a limited role of the arrestin-3-based scaffolding in the control of the JNK signaling. However, JNK family kinases are activated by many other MAP3Ks (Johnson, 2011; Peterson et al., 2022; Castro-Torres et al., 2024; Mordente et al., 2024). Here we demonstrated that arrestin-3 is a versatile scaffold that binds multiple MAP3Ks, including both isoforms of ZAK, MEKK1/2/4, and TAK1, and facilitates JNK3 activation triggered by these kinases (**Fig. 1**). Our discovery that arrestin-3 interacts with members of all three major MAP3K families (MEKK, MLK, and ASK) suggests that it possesses a versatile kinase-docking surface. This allows arrestin-3 to facilitate signaling by multiple MAP3Ks. This is significant from the mechanistic standpoint, because arrestin-3 scaffolds JNK activation cascade in its basal conformation, in contrast to the ERK pathway, which is assembled by GPCR-bound arrestins (Zheng et al., 2023; Gurevich and Gurevich, 2024). The observation that co-expression of arrestin-3 increases the concentration of JNK3 in cells – the effect it does not exert on MKK4/7 or MAP3Ks – suggests that a proportion of free arrestin-3 in the cytosol exists as a pre-assembled complex with JNK3. Apparently, the MAP3Ks activated in response to various stimuli provide signaling input, thereby making the arrestin-3-based JNK activation scaffold productive.

In HEK293-derived AKO cells lacking both non-visual arrestins, we identified ZAK, the two isoforms of which are particularly abundant in this cell type (Figs. 1J, S3, S4), as the major player in arrestin-3-assisted JNK3 activation (**Figs. 2, 4**), as well as in spontaneous and stressor-induced cell death (**Fig. 6**). This is consistent with previous observations that arrestin-3 facilitates JNK3α2 activation in the absence of ASK1 (Perry-Hauser et al., 2022). Our findings highlight a previously unappreciated role of ZAK as an initiator of arrestin-3-facilitated signaling to JNK3 in certain cells. We showed that in AKO cells arrestin-3-dependent enhancement of ZAK signaling plays a key role in both acute stress response to drugs and in the viability of unstressed cells (**Figs. 6, S11**). Our finding that ZAK KO cells demonstrate lower spontaneous death than cells endogenously expressing ZAK suggests that ZAK-initiated signaling sets a threshold for cell survival in this line. Importantly, our data with ZAK KO cells demonstrate that without the activity of an upstream MAP3K the signaling cannot ensue regardless of the presence of downstream components of the cascade and their interactions with the scaffold. Arrestin-3 amplifies ZAK-induced pro-death signaling, whereas in the absence of ZAK the amplifier has no input, which results in increased cell survival (**Fig. 6**). ZAKα acts as a sensor of ribosomal stalling and metabolic stress (Wang et al., 2005; Vind et al., 2020; Wu et al., 2020; Snieckute et al., 2022). Our data suggest that arrestin-3 facilitates the translation of the perturbations sensed by ZAK to the cell death machinery involving JNK3. The ability of arrestin-3 to scaffold JNK3-activating cascades initiated by several MAP3Ks (**Fig. 1**) suggests that arrestin-3 likely also amplifies the effects of other stressors sensed by different MAP3Ks.

Our findings underscore the importance of elucidating the complement of endogenous MAP3Ks in different cell types. We analyzed four different cell lines and found that relative abundance of endogenous MAP3Ks in each is distinct (**Figs. 1J, S3, S4**). Available complement of endogenous MAP3Ks likely determines the sensitivity of each cell type to extra– and intra-cellular signals, including those inducing stress.

The data on the movements of MAP3Ks within the cell are sparse. The best-known example is c-Raf, which is activated upon its translocation to the plasma membrane due to binding GTP-liganded small G protein Ras (reviewed in (Terrell and Morrison, 2019)). The other examples include the translocation of MEKK1 (Alapati et al., 2014) and DLK (Wallbach et al., 2016) from the cytoplasm to the nucleus, as well as retrograde transport of palmitoylated DLK attached to vesicles in axons (Holland et al., 2016). Here we described an earlier unappreciated movement of ZAKα under stress: upon treatment with vincristine or anisomycin intracellular distribution of both ZAKα changed from fairly even throughout the cytoplasm to distinct cytoplasmic areas where it was concentrated (**Fig. 5**). Importantly, intracellular redistribution of arrestin-3 depends on ZAKα: subcellular localization of arrestin-3 expressed alone does not change upon anisomycin or vincristine treatment (Fig. 5). Concerted translocation of stress-responsive MAP3K ZAKα and arrestin-3 to the same subcellular locations supports the idea of their functional cooperation in the cell. It is tempting to speculate that this redistribution of signal-initiator MAP3K and scaffold protein that facilitates the transduction of its signal to effector MAPKs serves to enhance the response. The molecular mechanism of this process and the nature of cytoplasmic regions to which these proteins translocate remain to be elucidated.

Another important finding is that the scaffolding function of arrestin-3 does not strictly require full-length protein. Peptide experiments (**Figs. 2, 3, 4, 6, S6, S7, S8**) suggest that the N-terminus of the arrestin-3 is the main kinase-binding element. We found that T16 peptide (arrestin-3 residues 9-24) functions as a mini-scaffold recruiting ZAK and facilitating JNK3 activation by ZAK (**Figs. 2, 4**) and several other MAP3Ks (Figs. 1, 3, S6, S7) and increasing pro-death signaling initiated by ZAK under stress (**Fig. 6**). These data challenge the view that large multi-domain protein scaffolds are necessary to properly orient kinases of different levels to ensure efficient signal propagation in MAPK activation cascades (Brown and Sacks, 2009). It appears that a relatively small docking interface can bring MAP3K, MAP2K, and MAPK into proximity for effective signal transmission. Importantly, arrestin-3-derived peptides demonstrate greater selectivity towards individual MAP3Ks than full-length protein (**Figs. 3C, S6, S7**). This paves the way to the use of different peptides to selectively facilitate JNK3 activation initiated by specific MAP3Ks. The shortest peptide that we showed to be active is T14, which is 14 residues long (**Figs. 2, 3, S8**). As the length of an average amino acid residue measured from amine to amine is ∼3.5Å, in fully extended conformation the length of this peptide would be ∼49Å. For comparison, the distances between the farthest tips of the two domains of full-length non-visual arrestins are not much greater, ∼70-75Å (Han et al., 2001; Milano et al., 2002; Zhan et al., 2011a). In contrast to full-length proteins that can only be introduced into cells by gene therapy, the introduction of peptides does not require gene transfer, thereby avoiding associated risks (discussed in (Tang and Xu, 2020; Cetin et al., 2024; Limon, 2024)). Therefore, in contrast to large proteins, peptides are a rapidly expanding class of therapeutic agents with low toxicity (reviewed in (Wang et al., 2022; Sharma et al., 2023)).

To summarize, we showed that arrestin-3 is a versatile scaffold of JNK-activating cascades that facilitates the propagation of signals initiated by several MAP3Ks. Whether arrestin-3 facilitates stress-induced signaling from the same and/or other MAP3Ks to p38 remains to be elucidated. As far as JNK3 is concerned, our data suggest that the MAP3K arrestin-3 predominantly assists by arrestin-3 largely depends on the relative expression levels of endogenous MAP3Ks in a particular cell type. We also demonstrated the translocation in response to two different stress signals of MAP3K ZAKα, which apparently drives the movement of co-expressed arrestin-3 to the same subcellular locations, consistent with the participation of both proteins in the same biological function. The ability of arrestin-3-derived peptide T16 to sensitize cells to chemotherapy drugs vincristine and doxorubicin and to facilitate pro-death signaling induced by these stressors makes T16, as well as other arrestin-3-derived JNK-activating peptides, a novel type of tools suitable for anti-cancer therapy.

## Methods

### Cell Lines and Culture Conditions

Wild-type HEK293 (human embryonic kidney) and HEK293 arrestin-2/3 knockout (AKO) cells (a generous gift of Dr. A. Inoue, Tohoku University, Japan; described in (Grundmann et al., 2018) were cultured in Dulbecco’s modified Eagle’s medium (DMEM) supplemented with 10% fetal bovine serum (FBS) and 1% penicillin/streptomycin. The ZAK knockout (ZAK KO) cell lines were generated on the AKO background (creating triple-null lines) as described below. Mouse neuroblastoma Neuro2a cells were cultured in DMEM containing 10% FBS and 1% penicillin/streptomycin. Human neuroblastoma LAN5 cells were cultured in RPMI-1640 medium supplemented with 10% FBS, 1% penicillin/streptomycin, and 1% Non-Essential Amino Acids (NEAA). Human neuroblastoma SMS-KCN cells were cultured in Iscove’s Modified Dulbecco’s Medium (IMDM) containing 20% FBS, 1% penicillin/streptomycin, and 1x ITS-G supplement (GIBCO). All cells were maintained at 37°C in a humidified 5% CO₂ incubator. All cell lines used in this study were routinely tested for mycoplasma contamination using MycoDect Mycoplasma Detection Kit (Alstem, #MD200) and were confirmed to be negative. Cells were used at low passage numbers (typically fewer than 15 passages after thawing) to maintain phenotypic consistency.

### Generation of ZAK Knockout Cell Lines

ZAK knockout was performed in AKO cells. Guide RNAs (gRNAs) targeting exon 2 and exon 20 of the human *ZAK* gene were designed and cloned into the pSpCas9(BB)-2A-Puro (PX459) V2 vector (Addgene #62988). The following gRNA sequences were used: for targeting Exon 2: 5′-CACTATAGGTAAACCGAGCTGCCAC –3′ (forward) and 5’-CAGCTCGGTTTACCTATAGTGCAAA –3’ (reverse) (based on (Vind et al., 2020)); for targeting Exon 20: 5’-TCAAGCCTAGTTATGATCAC-3’ (forward) and 5’-GTGATCATAACTAGGCTTGA-3’ (reverse).

AKO cells were transfected with the plasmid mixture. 48 hours post-transfection, cells were selected with puromycin (2 µg/mL) for 2 days. Surviving single cells (by limiting dilution) were placed into 96-well plates and expanded for 3 weeks. Clones were validated by PCR genotyping and western blotting with an anti-ZAK antibody. Two independent clones (KO1 and KO2) confirmed to lack both ZAKα and ZAKβ proteins were selected for subsequent experiments. Two sets of PCR primers were used for validation. For WT (Exon 20 deletion site): 5’-gtgaaccagtccagaagctcg-3’ (forward) and 5’-cactggctcttgagtcagtctca-3’ (reverse). Expected fragments: 385bp for WT, none for KO. For KO confirmation (amplification across deletion): 5’-ggatgagatgagtgaggtacctgc –3’ (reverse) and 5’-gagatgtcgtctctcggtgc –3’ (forward). Expected fragment: 528bp for KO, none for WT.

### Plasmids and Transfection

Expression constructs encoding N-terminally Venus-tagged arrestin-3 and HA-ASK1 were described earlier (Breitman et al., 2012); HA-tagged MAP3Ks (ZAKα, ZAKβ, MEKK1, MEKK2, MEKK4, TAK1) were obtained from Addgene (#12182, 12187, 21632, 44160, 141193, 141195, respectively). Venus-tagged arrestin-3 peptides (T1A, T16, T15, T14) were generated by subcloning the corresponding cDNA fragments (encoding residues 1-25, 9-24, 7-21, and 11-24, respectively) into the pcDNA3 vector in-frame with Venus, as described (Zhan et al., 2016; Perry-Hauser et al., 2022). For all signaling and biochemical assays, cells were plated at 70–80% confluency 24 h before transfection. Transfections were performed with jetOPTIMUS reagent (Polyplus) according to the manufacturer’s instructions.

### JNK3 Activation Assays

To assess JNK3 activation, cells were co-transfected with HA-JNK3α2, the indicated MAP3Ks, and either Venus (control), Venus-arrestin-3, or Venus-peptide constructs. 48 hours post-transfection, cells were harvested in lysis buffer. The levels of phosphorylated JNK3 were determined by western blotting using a phospho-specific SAPK/JNK antibody (Thr183/Tyr185) at 1:1000 dilution (Cell signaling #4668). Phosphorylated JNK signals were normalized to the total HA-JNK3α2 or to loading control (GAPDH).

### Co-immunoprecipitation (Co-IP)

AKO cells were co-transfected with Venus-tagged constructs (arrestin-3, T16, T14) and HA-tagged MAP3Ks. Cells were lysed in buffer containing 25 mM HEPES (pH 7.3), 150 mM NaCl, 10% glycerol, 2 mM EDTA, 20 mM NaF, 0.5% NP-40, 1 mM Na_3_VO_4_, 2 mM benzamidine, 2 mM PMSF, and protease inhibitor cocktail (Sigma, P8340) (IP buffer). Lysates were clarified by centrifugation at 14,000 × g for 10 min at 4°C. Clarified lysates containing 500 μg of total protein were incubated with 20 μl of GFP-Trap magnetic beads (Chromotek, gtma-20) for 2 hours at 4°C with rotation. Beads were washed three times with 500 μl of IP buffer. Bound proteins were eluted in 100 μl of 2x Laemmli sample buffer at 95°C for 5 min and analyzed by western blotting.

### Western blotting

Protein samples were resolved by SDS-PAGE and transferred to PVDF membranes. Membranes were blocked in 5% non-fat dry milk and incubated with primary antibodies overnight at 4°C. The following antibodies were used at 1:1000 dilution: anti-HA (Cell signaling Cat#3724), anti-GFP (Takara; Cat#632381), anti-ZAK (ABclonal Biotechnology; Cat#A7371), anti-ASK1 (ABclonal; Cat#A12458), anti-MEKK1 (ABclonal; Cat#A16057), anti-MEKK2 (Cell signaling Technology; Cat#19607), anti-MEKK4 (ABclonal; Cat#A20874), anti-TAK1 (Cell signaling Technology: Cat#4505), anti-p-SAPK/JNK (Thr183/Tyr185) (Cell Signaling Technology: #4668), or anti-GAPDH (Cell signaling Technology: Cat#2118). After three 5 min washes HRP-conjugated secondary antibodies (Jackson ImmunoResearch: #115005003) were added for 1 h at room temperature, followed by three 5 min washes. The bands were detected by 2 min incubation with enhanced chemiluminescence reagent (ECL, Pierce #32209). Band intensities were quantified using ImageStudio software (LICORbio).

### Drug treatment

Cells were treated with the following agents prior to lysis: anisomycin: 300 nM for 1 h (ribotoxic stress); vincristine: 100 nM for 24 h (microtubule destabilization); doxorubicin: 20 nM for 24 h (DNA damage). Vehicle controls (DMSO) were included in all experiments.

### Confocal microscopy and image analysis

AKO and ZAK KO cells were plated on fibronectin-coated glass coverslips and co-transfected with Venus-arrestin-3 and mCherry-ZAKα. 24 h post transfection, cells were transferred to poly-D-lysine (Sigma P6407, 50 μg/ml for 2 h) coated MatTek glass-bottom dishes. Cells were treated with vehicle (DMSO), anisomycin (300 nM), or vincristine (500 nM) for 16 hours. Nuclei were stained with NucBlue Live ReadyProbes (ThermoFisher; Cat#R37605) prior to imaging. Live-cell images were acquired using a Nikon A1r resonant scanning Eclipse Ti2 HD25 confocal microscope with a 10x (Nikon #MRD00105, CFI60 Plan Apochromat Lambda, N.A. 0.45), 20x (Nikon #MRD00205, CFI60 Plan Apochromat Lambda, N.A. 0.75), and 60x (Nikon #MRD01605, CFI60 Plan Apochromat Lambda, N.A. 1.4) objectives operated by NIS-24 Elements AR v4.5 acquisition software. Growth conditions were maintained (37°C, 5% CO_2_, 90% humidity) using a Tokai Hit stage incubator (Tokai Hit STXG Incubation System).

### Image Analysis

Fluorescence line scans were generated using the “Plot Profile” function in ImageJ/Fiji. To quantify distribution of arrestin-3 and ZAKα, we calculated an Enrichment Index as previously described (Varandas et al., 2016; Ripoll et al., 2024). Briefly, regions of interest corresponding to the whole cell were defined to measure the mean fluorescence intensity of the total cell. High-intensity areas were segmented using Otsu thresholding, and the mean intensity of these enriched regions was measured. The Enrichment Index was defined as the ratio of the mean intensity of these areas to the mean intensity of the whole cell.

### Flow cytometry

AKO or AKO + ZAK KO cells were co-transfected with HA-JNK3α2 and Venus, Venus-arrestin-3, or Venus-T16. For drug-induced death: 36 hours post-transfection, cells were exposed to anisomycin (1.5μM or 3μM) or DMSO (control) for 48 hours. For spontaneous death cells were harvested at 24 hours or 48 hours post-transfection without drug treatment.

Cells were harvested by centrifugation (500 x g, 5 min), washed in DPBS, and stained with LIVE/DEAD Fixable Far Red Dead Cell Stain (Invitrogen, L34973) according to manufacturer’s instructions. Cells were then fixed in 3.7% paraformaldehyde (Electron Microscope Sciences, 15714) in DPBS and filtered through 40 μm cell strainers (Fisher, 22363547). Flow cytometric analysis was performed immediately using a BD Fortessa flow cytometer (BD Biosciences).

Analysis was performed using FlowJo software (BD Biosciences). Cells were distinguished from debris using forward and side scatter parameters (FSC-A and SSC-A). Transfected cells were identified by FITC fluorescence, with gates established using unstained control populations.

### Statistical Analysis

Data are presented as means ± standard error of the mean (SEM) from at least three independent biological replicates (N≥3). For image and flow cytometry analyses, investigators were blinded to the group allocation during data collection and quantification. Statistical analyses were performed using GraphPad Prism software (Dotmatics). Comparisons between two groups were analyzed using an unpaired two-tailed Student’s t-test. For comparisons between two treatments across multiple independent conditions, multiple unpaired t-tests were performed with the False Discovery Rate (FDR) set at 5% using the two-stage step-up method of Benjamini, Krieger, and Yekutieli. Multiple group comparisons were analyzed using one-way ANOVA with Dunnett’s correction for multiple comparisons. When the assumption of equal variances was not met, Brown-Forsythe and Welch ANOVA tests were used with Dunnett’s correction for multiple comparisons. Experiments involving two variables (e.g., Genotype × Treatment, or Peptide × Kinase) were analyzed using two-way ANOVA with Sidak’s correction for multiple comparisons. Time-course experiments involving three variables (Time × Genotype × Transfection) were analyzed using three-way ANOVA. A p value (or FDR-adjusted q value) of < 0.05 was considered statistically significant. Statistical significance is indicated, as follows: *, p<0.05; **p<0.01; ***p<0.001; ****p<0.0001. In some figures p values are shown.

## Acknowledgements

Supported by NIH RO1 EY011500 and Cornelius Vanderbilt Endowed Chair (Vanderbilt University) (VVG). The authors thank Dr. Oleg Kovtun (Vanderbilt University) for his expert advice and insightful discussions regarding image quantification strategies.

## Data Availability

The data that support the findings of this study are available within the paper and its Supplementary Information. Uncropped and unprocessed scans of all western blots are provided in the Source Data file. Raw data and materials are available from the corresponding author upon request.

## Author contributions

C.Z., V.V.G, and E.V.G. conceived the experiments. C.Z. and E.V.G performed the experiments. C.Z., V.V.G, R.S., and E.V.G. wrote the manuscript with contributions from all authors. All coauthors contributed to discussions of the protocol and results. E.V.G. and V.V.G. supervised the project.

## Footnote

^1^We use systematic names of arrestin proteins, where the number after the dash indicates the order of cloning: arrestin-1 (historic names S-antigen, 48 kDa protein, visual or rod arrestin), arrestin-2 (β-arrestin or β-arrestin1), arrestin-3 (β-arrestin2 or hTHY-ARRX), and arrestin-4 (cone or X-arrestin).

