## Supplementary material for "Arrestin-3 scaffolds multiple MAP3Ks driving stress-induced JNK3 activation and cell death": S1

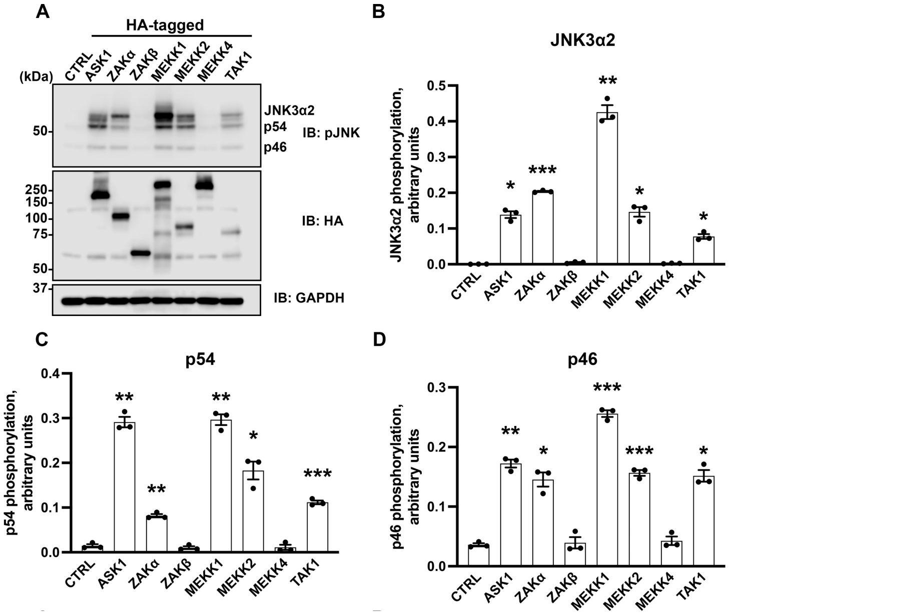


**Figure S1. Effects of MAP3Ks on JNK activation in AKO cells. A.** Western blot of AKO cells co-transfected with HA-JNK3α2 and indicated MAP3Ks. Upper panel: western blot with anti-ppJNK3 antibody. Lower panel: expression of indicated MAP3Ks was determined with anti-HA antibody (HA). The intensity of doubly phosphorylated JNK3α2 (B), endogenous p54 (C) and p46 (D) was quantified (N=3). Statistical significance of the differences with cells transfected with empty vector (CTRL) was determined by Welch's one-way ANOVA followed by Dunnett's correction for multiple comparisons. p values are indicated, as follows: *, $p<0.05$; **, $p<0.01$; ***, $p<0.001$.

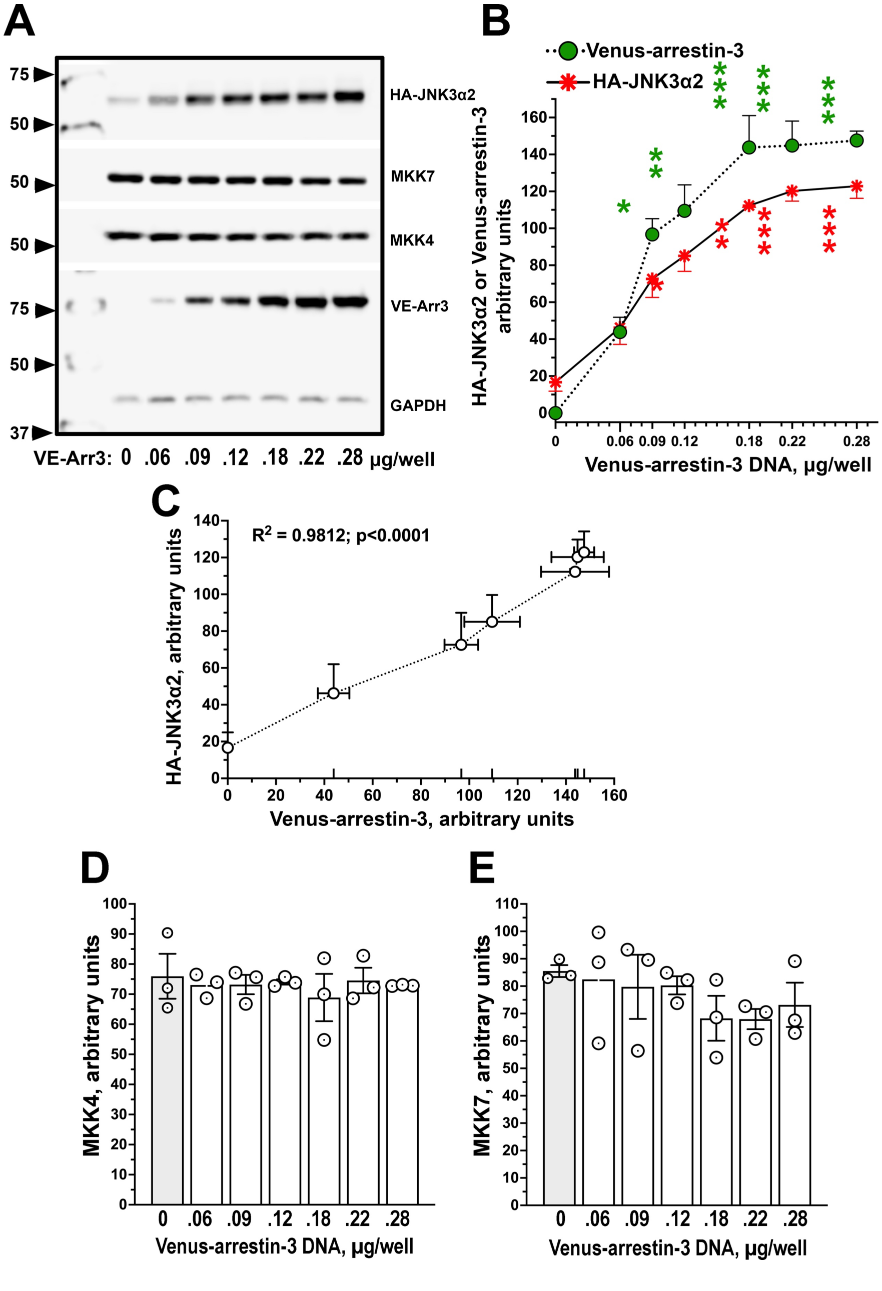

A
75
50
50
50
75
50
37
HA-JNK3α2
MKK7
MKK4
VE-Arr3
GAPDH
VE-Arr3: 0 .06 .09 .12 .18 .22 .28 μg/well
B
Venus-arrestin-3
HA-JNK3α2
HA-JNK3α2 or Venus-arrestin-3 arbitrary units
Venus-arrestin-3 DNA, μg/well
0 0.06 0.09 0.12 0.18 0.22 0.28
C
R2 = 0.9812; p<0.0001
HA-JNK3α2, arbitrary units
Venus-arrestin-3, arbitrary units
D
MKK4, arbitrary units
0 .06 .09 .12 .18 .22 .28
Venus-arrestin-3 DNA, μg/well
E
MKK7, arbitrary units
0 .06 .09 .12 .18 .22 .28
Venus-arrestin-3 DNA, μg/well

**Figure S2. Arrestin-3 increases the JNK3α2 expression. (A**) Representative Western blot showing the expression of HA-JNK3α2 detected with anti-HA antibody, endogenous MAP2Ks MKK7 and MKK4 detected with anti-MKK7 and anti-MKK4 antibodies, respectively, in the presence of different expression of Venus-arrestin-3 detected with anti-GFP antibody. **(B)** Quantification of the Western blot data for HA-JNK3α2 and Venus-arrestin-3. Significance of the differences with cells transfected with 0.06 μg of the plasmid encoding Venus-arrestin-3 was determined by one-way ANOVA and shown, as follows (green, Venus-arrestin-3; red – HA-JNK3α2): *, $p<0.05$; **, $p<0.01$; ***, $p<0.001$. **C.** Covariation of the expression of Venus-arrestin-3 and HA-JNK3α2 was analysed by Pearson XY correlation test; $R^2$ and p value indicated. **(D,E)** Quantification of the Western blot for MKK4 **(D)** and MKK7 **(E)**. The expression of endogenous MKK4 and MKK7 was not significantly affected by the expression of Venus-arrestin-3.

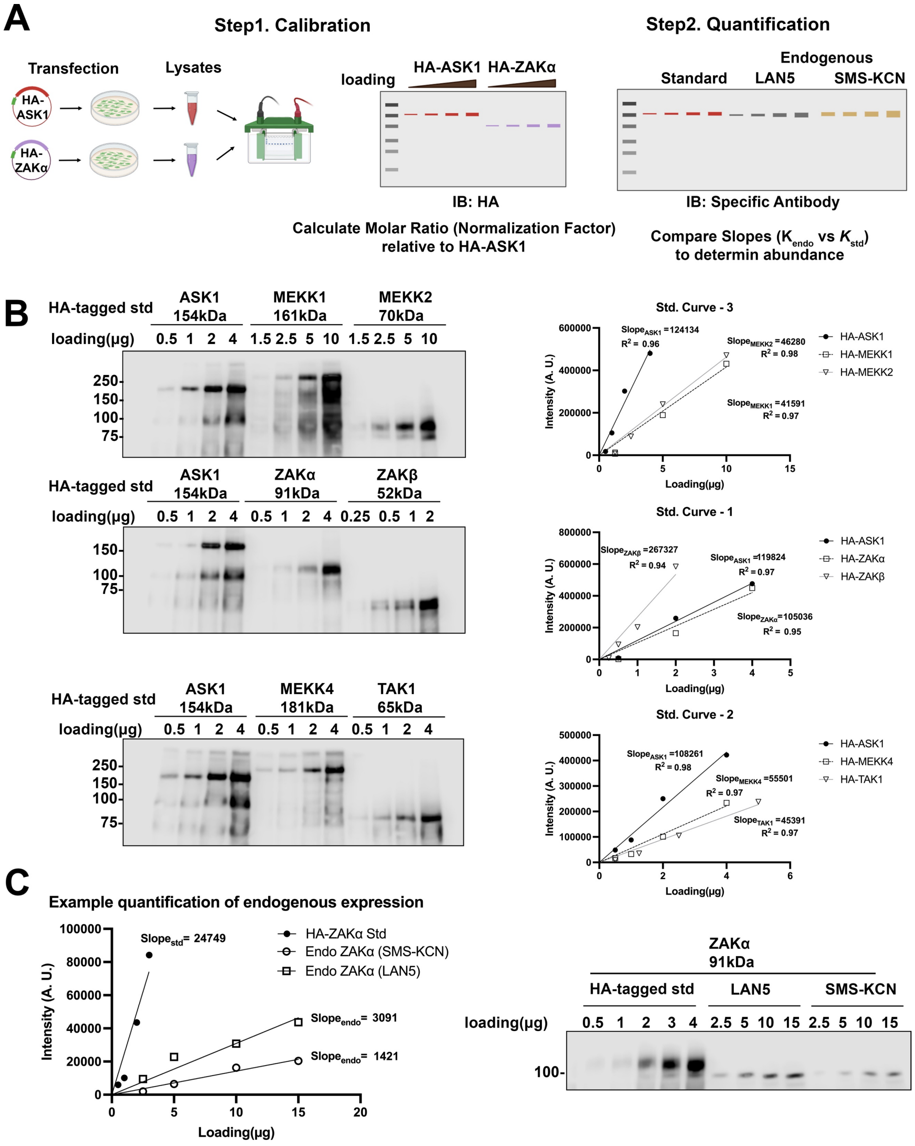


**Figure S3. Quantitative immunoblotting strategy for the relative abundance of endogenous MAP3Ks. A.** Schematic representation of the two-step absolute quantification workflow. Step 1: Calibration curves are established using increasing amounts of transfected HA-tagged MAP3Ks detected via an anti-HA antibody. Step 2: The relative expression of endogenous MAP3Ks across different cell lines is determined using MAP3K-specific antibodies, with the pre-quantified HA-tagged MAP3Ks serving as reference standards. **(B)** Representative western blots and corresponding calibration curves of the HA-tagged MAP3K reference standards. Signals were quantified to equalize the amounts of different HA-MAP3K standards prior to endogenous MAP3K quantification. **(C)** A representative example demonstrating the quantification workflow for ZAKα. The standard curve generated from the HA-ZAKα reference was utilized to determine the relative abundance of endogenous ZAKα in whole-cell lysates.

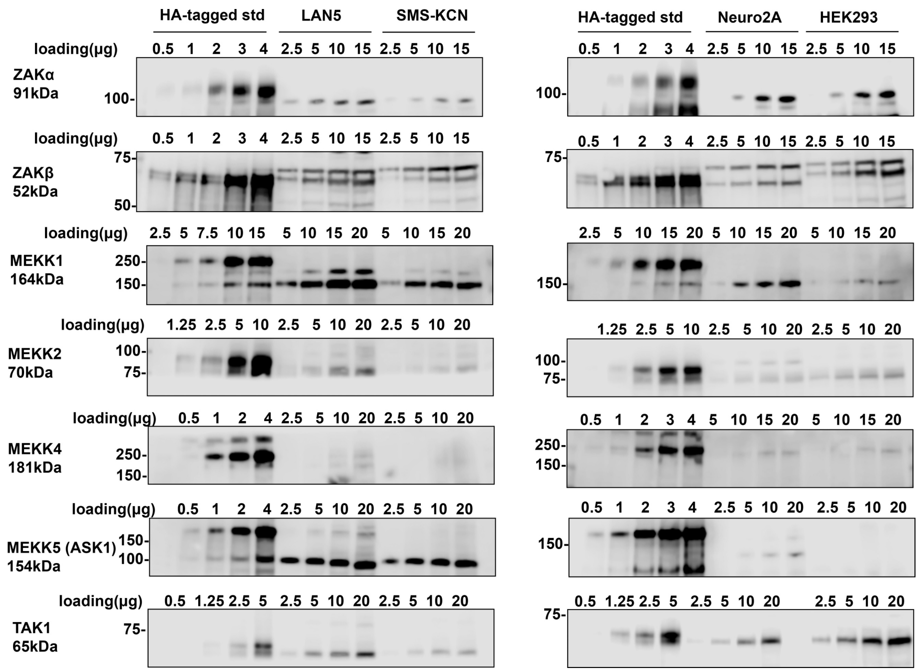


**Figure S4. Endogenous expression profiles of MAP3Ks in different cell lines.** Representative western blots showing the expression levels of the indicated endogenous MAP3Ks in diverse cellular contexts, including human neuroblastoma (LAN5, SMS-KCN), mouse neuroblastoma (Neuro2a), and HEK293 cell lysates. Blots were probed with kinase-specific antibodies. To enable accurate cross-blot normalization and relative quantification,

**Deleted:** Neuro2A

corresponding HA-tagged MAP3K reference standards (HA-tagged std) were loaded in parallel. The total protein loading amounts (μg) for each cellular lysate are indicated above respective lanes.

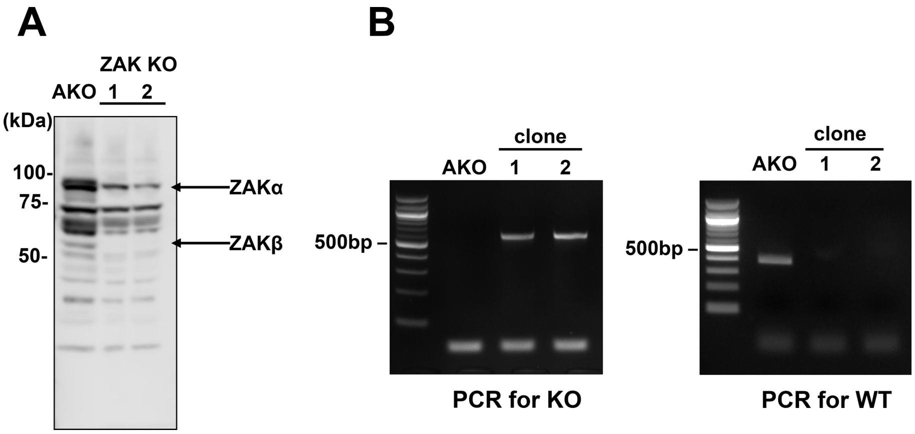


**Figure S5. Verification of ZAK knockout in HEK293 cells. A.** Western blot of lysates from parental (AKO) and two independent ZAK knockout clones (KO1, KO2). The membrane was probed with an anti-ZAK antibody that recognizes both α and β isoforms. **B.** Agarose gels for the analysis of ZAK knockout by PCR. The bands expected with WT ZAK gene and in correct knockout are 385 bp and 528 bp long, respectively.

**Deleted:** 528 bp and

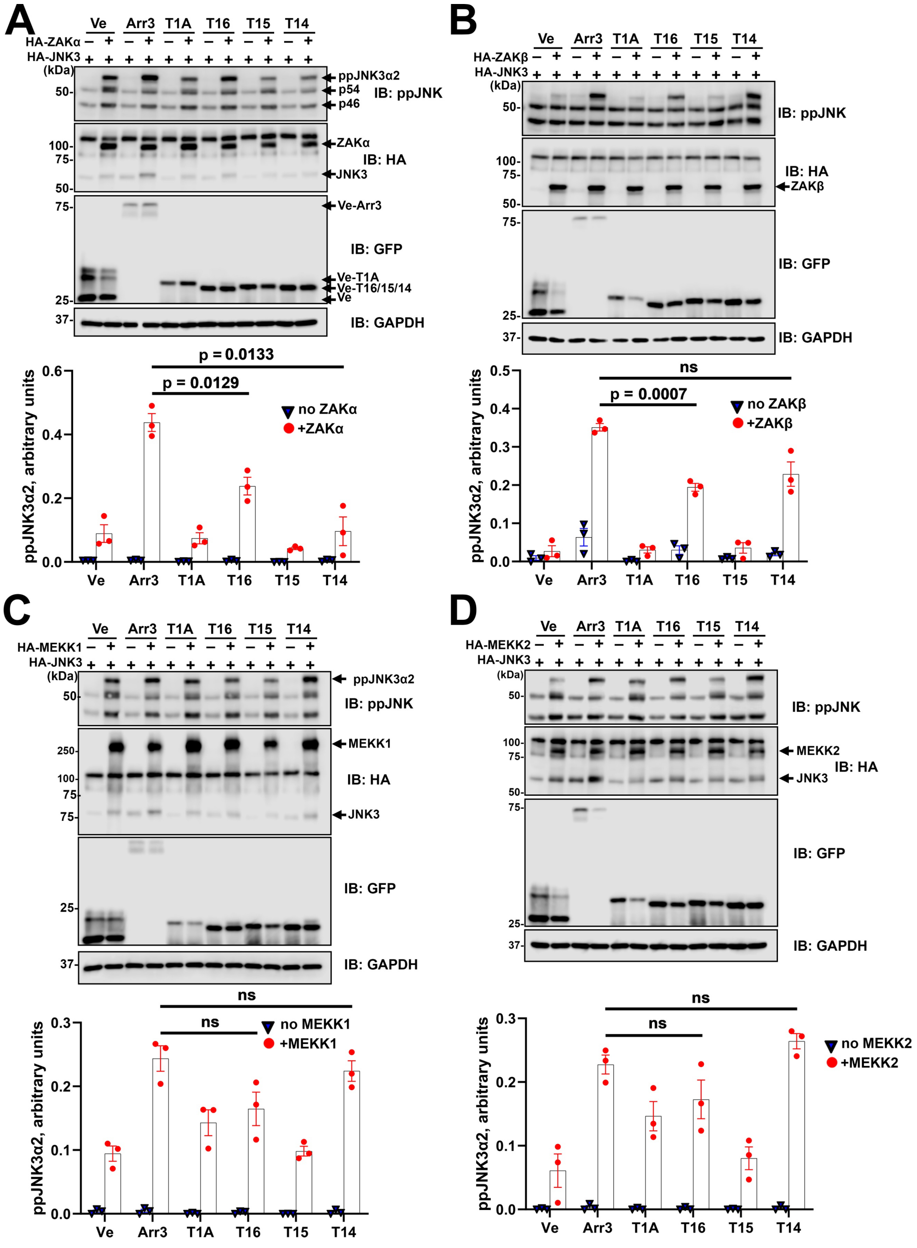
A
Ve
Arr3
T1A
T16
T15
T14
HA-ZAKα
HA-JNK3
(kDa)
ppJNK3α2
p54
p46
IB: ppJNK
ZAKα
IB: HA
JNK3
Ve-Arr3
IB: GFP
Ve-T1A
Ve-T16/15/14
Ve
IB: GAPDH
p = 0.0133
p = 0.0129
no ZAKα
+ZAKα
ppJNK3α2, arbitrary units
B
HA-ZAKβ
IB: ppJNK
IB: HA
ZAKβ
IB: GFP
IB: GAPDH
ns
p = 0.0007
no ZAKβ
+ZAKβ
C
HA-MEKK1
ppJNK3α2
IB: ppJNK
MEKK1
IB: HA
JNK3
IB: GFP
IB: GAPDH
ns
no MEKK1
+MEKK1
D
HA-MEKK2
IB: ppJNK
MEKK2
IB: HA
JNK3
IB: GFP
IB: GAPDH
ns
no MEKK2
+MEKK2

**Figure S6. Facilitation of JNK3 activation by arrestin-3-derived peptides. A.** ZAKα. **B.** ZAKβ. **C.** MEKK1. **D.** MEKK2. Representative western blots with anti-ppJNK, anti-HA, and anti-GFP antibodies are shown. GAPDH served as a loading control. Bar graphs: quantification of ppJNK3α2 (N=3). Statistical significance of the differences was determined in the MAP3K-transfected groups using Welch's one-way ANOVA with Dunnett's correction for multiple comparisons; p values are indicated.

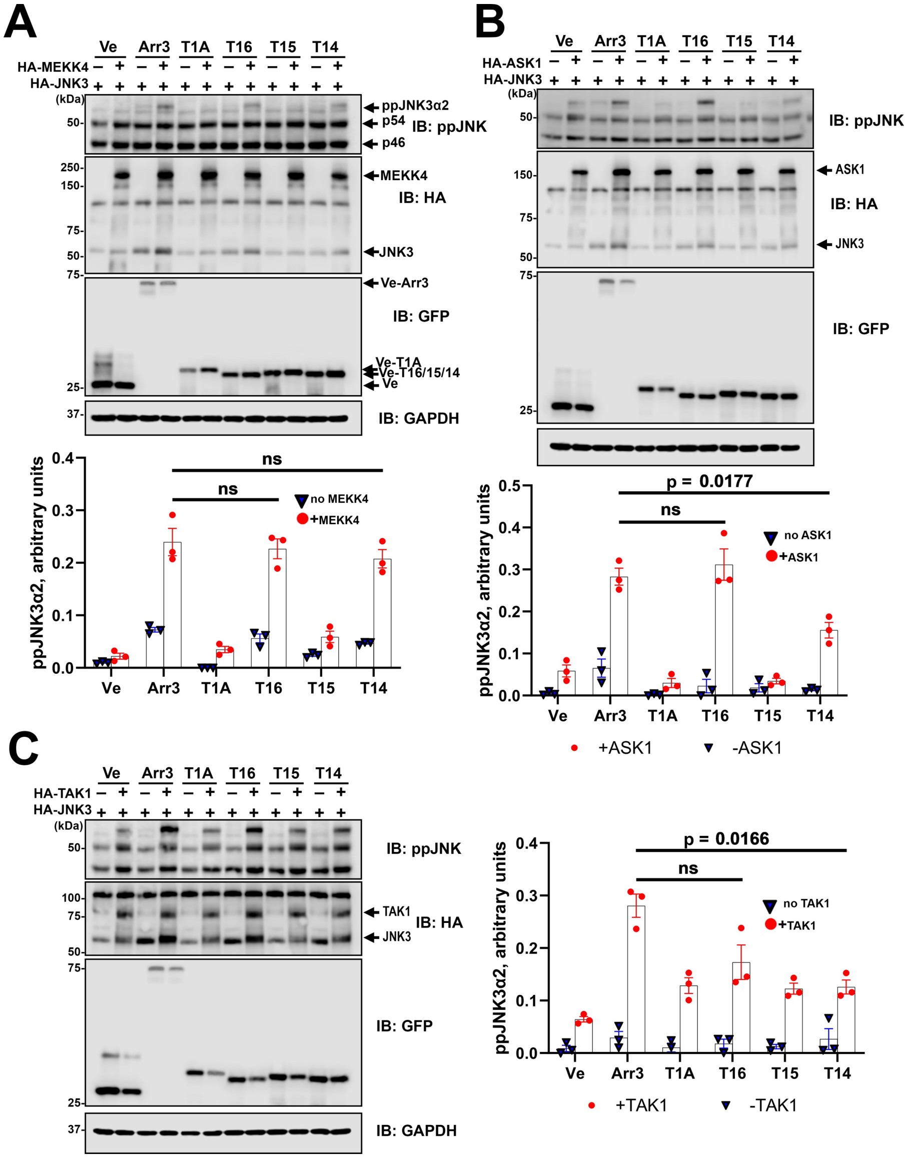


**Figure S7. Facilitation of JNK3 activation by arrestin-3-derived peptides. A.** MEKK4. **B.** ASK1. **C.** TAK1. Representative western blots with anti-ppJNK, anti-HA, and anti-GFP antibodies are shown. GAPDH served as a loading control. Bar graphs: quantification of ppJNK3α2 (N=3). Statistical significance of the differences was determined in the MAP3K-transfected groups using Welch's one-way ANOVA with Dunnett's correction for multiple comparisons; p values are indicated.

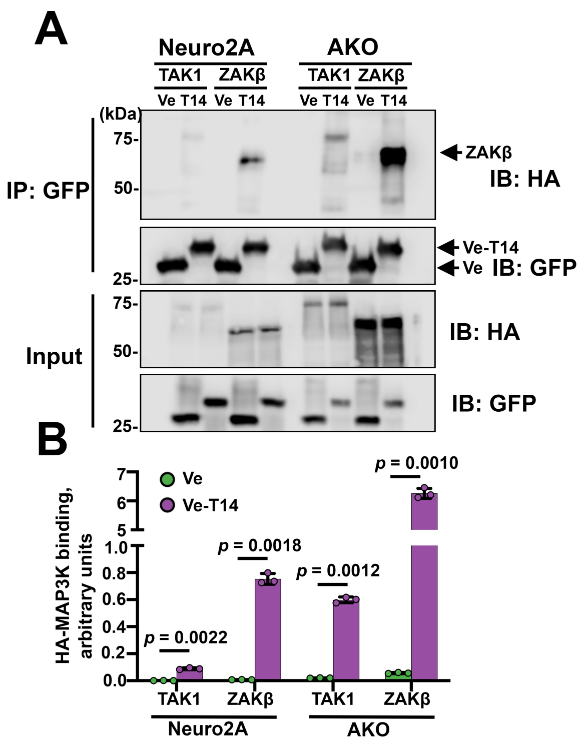


**Figure S8. Co-immunoprecipitation of T14 peptide with MAP3Ks. A.** Neuro2a and AKO cells were transfected with HA-TAK1 or HA-ZAKβ and Venus (control) or Venus-T14. Venus and Venus-T14 were immunoprecipitated with GFP-Trap beads. Western blot with anti-HA antibody revealed co-precipitated HA-TAK1 and HA-ZAKβ. **B.** Quantification of HA-MAP3K bands. Statistical significance of the differences between Venus and Venus-T14 was determined by multiple unpaired t-tests with FDR set at 5%; p values are indicated.

Deleted: Neuro2A

Deleted: a

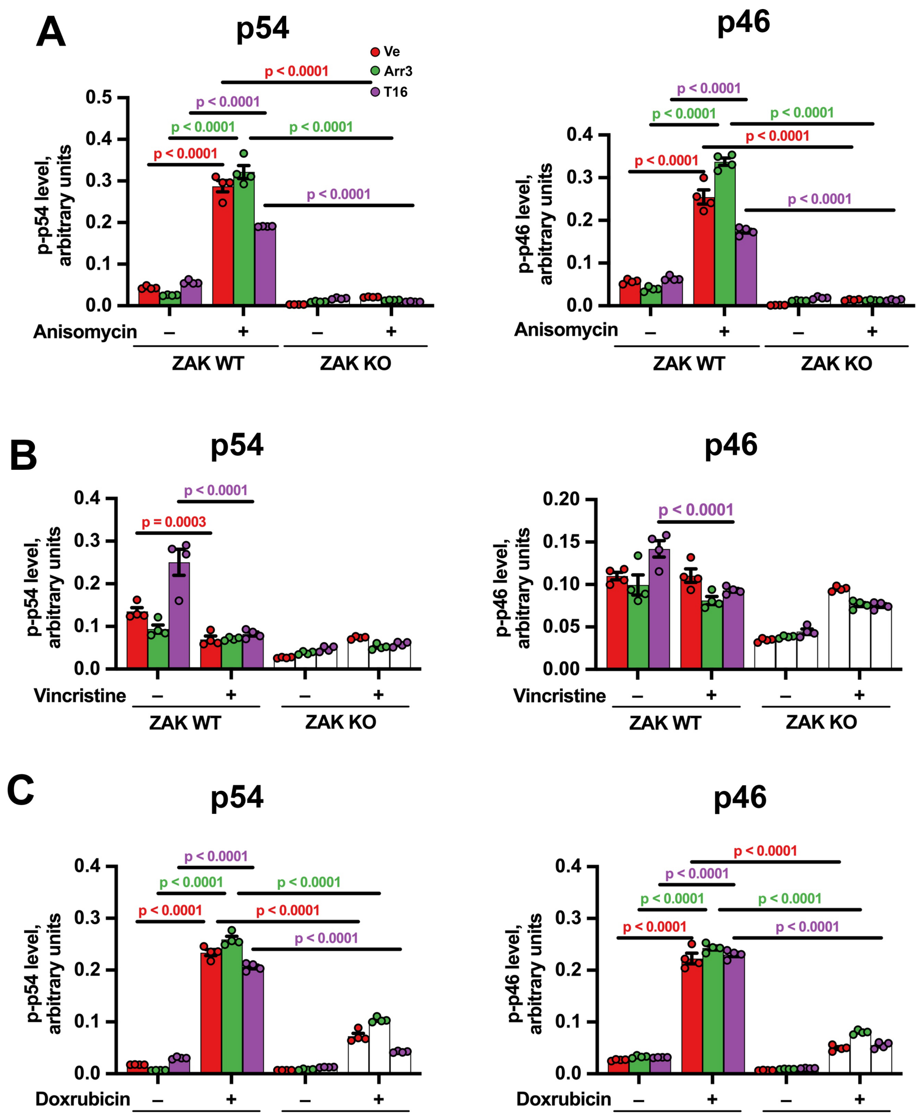


**Figure S9. The effect of stressors on the activation of endogenous JNKs.** The intensity of of p46 and p54 bands (endogenous JNK isoforms) detected by ppJNK antibody (representative blots are shown in Fig. 4) in cells treated with anisomycin (**A**), vincristine

**Deleted:** anysomycin

**Formatted:** Font: Bold

(**B**), and doxorubicin (**C**) was quantified. Statistical significance of the differences was determined by two-way ANOVA with Sidak's correction for multiple comparisons; p values are shown.

Formatted: Font: Bold

Formatted: Font: Bold

Deleted: by two

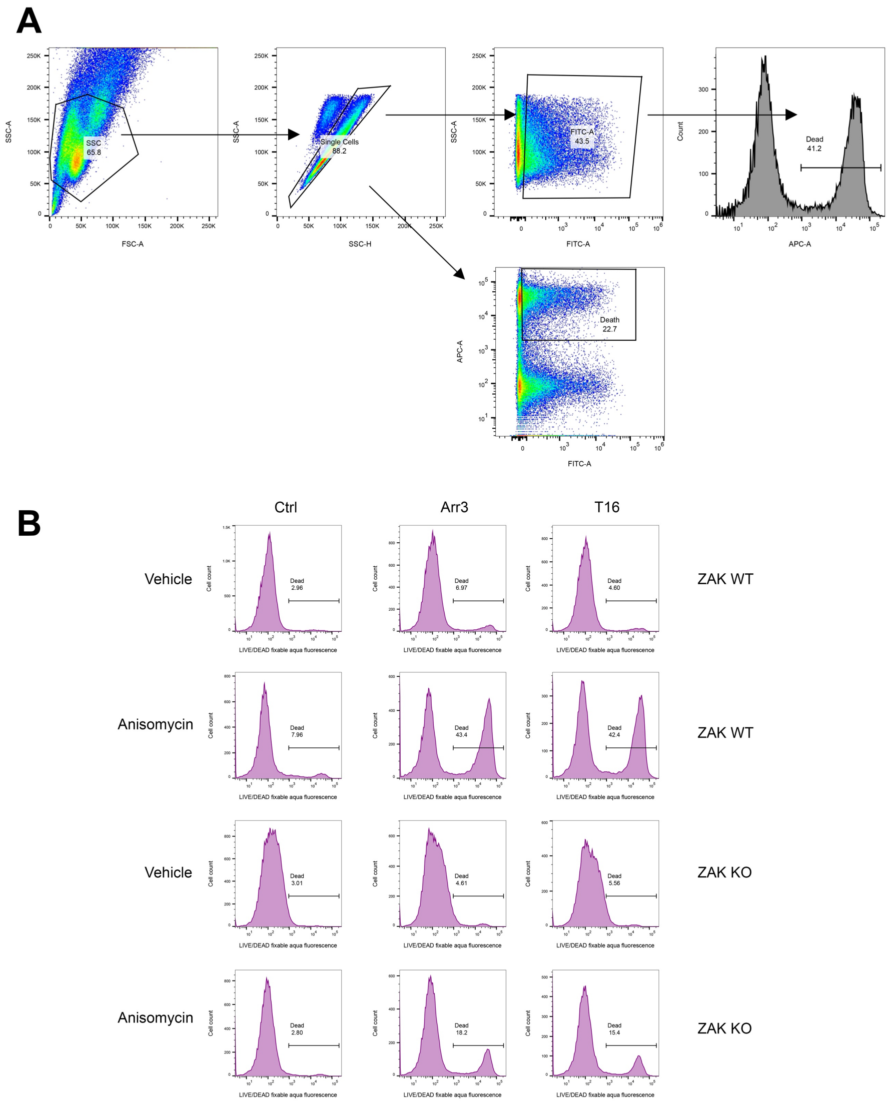


**Figure S10. Flow cytometry gating strategy and analysis of cell death. A.** Detailed gating strategy for flow cytometry analysis. Sequential gating shows: (1) Forward and side scatter to exclude debris, (2) Single cell discrimination, (3) Venus-positive (transfected) cell selection, and (4) LIVE/DEAD staining analysis. **B.** Complete flow cytometry data for all experimental conditions. ZAK WT and ZAK KO cells expressing Venus, Venus-arrestin-3, or Venus-T16 with HA-JNK3α2 were treated with DMSO (control) or anisomycin. Both scatter plots and histogram overlays are shown for comprehensive analysis of cell death responses.

**A.** Detailed sequential gating strategy used for all flow cytometry analyses. Used sequence was, as follows:

1. **FSC-A vs. SSC-A**: Exclusion of cellular debris.
2. **FSC-H vs. FSC-A**: Doublet discrimination to select single cells.
3. **FITC-A (Venus)**: Gating for Venus-positive cells to analyze only transfected cells expressing Venus control, Venus-arrestin-3, or Venus-T16.
4. **APC-A (LIVE/DEAD)**: Analysis of cell death within the Venus-positive population using LIVE/DEAD™ Fixable Far Red Dead Cell Stain.

**B.** Representative flow cytometry histograms for the cell death analysis shown in Figure 6. Histograms show the distribution of dead cells (APC-A signal) in ZAK WT and ZAK KO cells expressing Venus, Venus-arrestin-3, or Venus-T16, treated with DMSO (control) or anisomycin. These histograms complement the density plots shown in Figure 6D.

**Deleted:** B

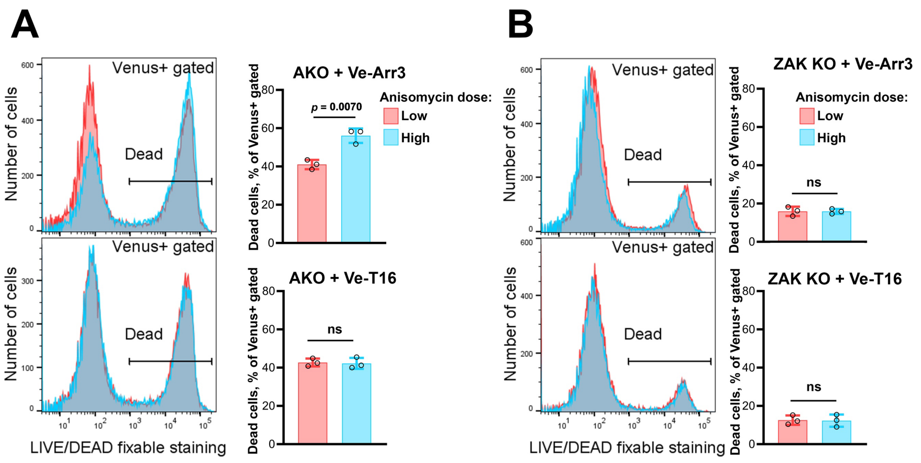

**Figure S11. The effect of arrestin-3, but not of T16, depends on the dose of anisomycin.** AKO (**A**) and AKO + ZAK KO (**B**) cells were treated with 1.5 (low dose) or 3 μM (high dose) of anisomycin. Cells were stained with LIVE/DEAD™ Fixable Far Red Dead Cell Stain, analyzed, as described in Methods, and the fraction of dead cells was quantified. Statistical significance of the differences was determined by Welch's t test; significant p value is shown.

**Deleted:** sorted
